# AlaRS-mediated lactylation shields PARP1 from ubiquitination to promote HNSCC progression

**DOI:** 10.64898/2026.05.18.725782

**Authors:** Jingjing Zhang, Xinru Meng, Qianni Qin, Qikai Zhou, Ying Lu, Runping Qin, Yujuan Yan, Chang Liu, Siwei Zhang, Xiaoyun Luo, Gen Liu, Yingjie Bian, Zhong-Wei Zhou, Ji-Chang Zhou, Jinjun Gao, Lanlan Wei, Bin Ma, Paul Schimmel, Litao Sun

## Abstract

Metabolic reprogramming in solid tumors causes massive lactate accumulation, yet how this drives oncogenesis remains incompletely understood. Here, we identify alanyl-tRNA synthetase (AlaRS) as a cellular lactyltransferase that promotes head and neck squamous cell carcinoma (HNSCC) progression. AlaRS directly catalyzes site-specific lactylation of the DNA repair protein PARP1 at lysines K249 and K667. Crucially, we uncover a competitive post-translational crosstalk wherein this lactylation directly antagonizes PARP1 ubiquitination. This "lactyl-shield" prevents proteasomal degradation, hyper-stabilizing the PARP1-Mortalin complex and sustaining tumor proliferation via p53/p21 dysregulation. To therapeutically exploit this mechanism, we identified chelerythrine (CHE) as a potent, selective inhibitor that directly binds the AlaRS catalytic center. CHE abrogates AlaRS lactyltransferase activity, destabilizes PARP1, and robustly suppresses HNSCC xenograft growth in vivo. These findings establish a novel metabolic-post-translational axis linking lactate accumulation to oncoprotein stabilization, providing a blueprint for targeting tRNA synthetase moonlighting functions in cancer.

## 1. Introduction

Head and neck squamous cell carcinoma (HNSCC), the sixth most common malignancy worldwide, is characterized by high metabolic activity, rapid proliferation, and poor clinical outcomes. Despite advances in multimodal treatment strategies, including radiation therapy and chemotherapy, the five-year overall survival rate remains below 50%, with more than 65% of patients experiencing recurrence, metastasis, or both^1–3^. This persistent therapeutic challenge highlights the urgent need to elucidate the specific molecular drivers of HNSCC pathogenesis and to identify novel therapeutic targets.

Metabolic reprogramming, a hallmark of cancer, enables tumor cells to meet their increased bioenergetic and biosynthetic demands in support of proliferation, invasion, and metastasis^4, 5^. This adaptive process is dynamic, evolving throughout cancer progression^6, 7^. Elucidating the mechanisms by which metabolic reprogramming promotes tumor development offers the potential to selectively target these pathways and suppress associated malignant phenotypes. Importantly, post-translational modifications (PTMs) form a critical bridge between metabolic reprogramming and protein functional regulation, as numerous metabolites serve as direct substrates for these modifications^8, 9^. Among them, lactate has been identified as the substrate for lysine lactylation, establishing a direct link between glycolytic metabolism and protein function^10, 11^. Histone lactylation, notably at H3K18, has been implicated in various cancer-related processes including tumor initiation^12, 13^, progression^14, 15^, and immune evasion^6^. Beyond histones, lactylation also regulates non-histone functions. For instance, lactylation of methyltransferase 16 (METTL16) in gastric cancer enhances m^6^A modification of *ferredoxin 1* (*FDX1*) mRNA and promotes elesclomol efficacy^16^. Given that HNSCC exhibits high glycolytic flux and lactate accumulation^11, 17^, defining the specific targets and functional impact of lactylation in this context is essential for understanding disease mechanisms and revealing new therapeutic opportunities.

Cancer cells exhibit an elevated demand for protein synthesis to sustain rapid proliferation, a process supported by the increased expression and functional versatility of aminoacyl-tRNA synthetases (aaRSs)^18^. This ancient enzyme family comprise 20 members, each specific to an amino acid, responsible for ligating amino acids to their corresponding tRNAs during protein synthesis. Beyond their canonical roles, aaRSs exhibit moonlighting functions in cancer, contributing to network dysregulation through non-canonical activities^19–21^. AaRSs dysregulation is now recognized as a driver of tumorigenesis and metastasis via diverse mechanisms, including translational reprogramming, metabolite sensing, and PTM catalysis^22–24^. Given their pivotal roles in cancer, aaRSs-specific compounds are gaining attention as potential therapeutic agents. For example, LysRS interacts with the 67-kDa laminin receptor to promote metastasis in lung and breast cancers, an interaction selectively disrupted by YH16899 without impairing aminoacylation^25^. Similarly, overexpressed LeuRS activates mTORC1 via RagD binding, and BC-LI-0186 inhibits this interaction to suppress signaling ^26, 27^.

Here, we identify AlaRS as a previously unrecognized lactyltransferase in HNSCC that operates beyond its canonical role in translation. We show that AlaRS catalyzes site-specific lactylation of PARP1 at lysine residues shared with ubiquitination sites, competitively blocking ubiquitin-proteasome degradation, thereby stabilizing PARP1 and sustaining oncogenic signaling through dysregulation of the p53/p21 axis. Targeting this axis, we identify chelerythrine (CHE) as a direct inhibitor of AlaRS lactyltransferase activity that suppresses HNSCC growth *in vitro* and *in vivo*. These findings establish a metabolic and post-translational regulatory axis centered on AlaRS that links lactate accumulation to oncoprotein stabilization, offering a novel therapeutic strategy for HNSCC.

## 2. Results

### 2.1. AlaRS is upregulated in HNSCC and correlates with poor prognosis

To systematically evaluate the involvement of aaRSs in HNSCC, we first analyzed mRNA expression levels of all 20 aaRSs in HNSCC tumors compared to normal tissue samples from The Cancer Genome Atlas (TCGA) dataset (**Figure 1A**). *TrpRS*, *AlaRS*, and *CysRS* exhibited significant upregulation in tumors. Proteomic analysis from the Clinical Proteomic Tumor Analysis Consortium (CPTAC) (**Figure 1B**) and Venn diagram analysis further validated the upregulation of TrpRS and AlaRS (**Figure 1C**). Notably, Kaplan-Meier analysis revealed that high AlaRS expression strongly correlated with shorter overall survival (**Figure 1D**), whereas TrpRS expression showed no prognostic association (**Figure S1**). These findings collectively suggest that AlaRS is upregulated at both the transcriptional and protein levels in HNSCC, and its high expression correlates with poor survival.

**Figure 1.**
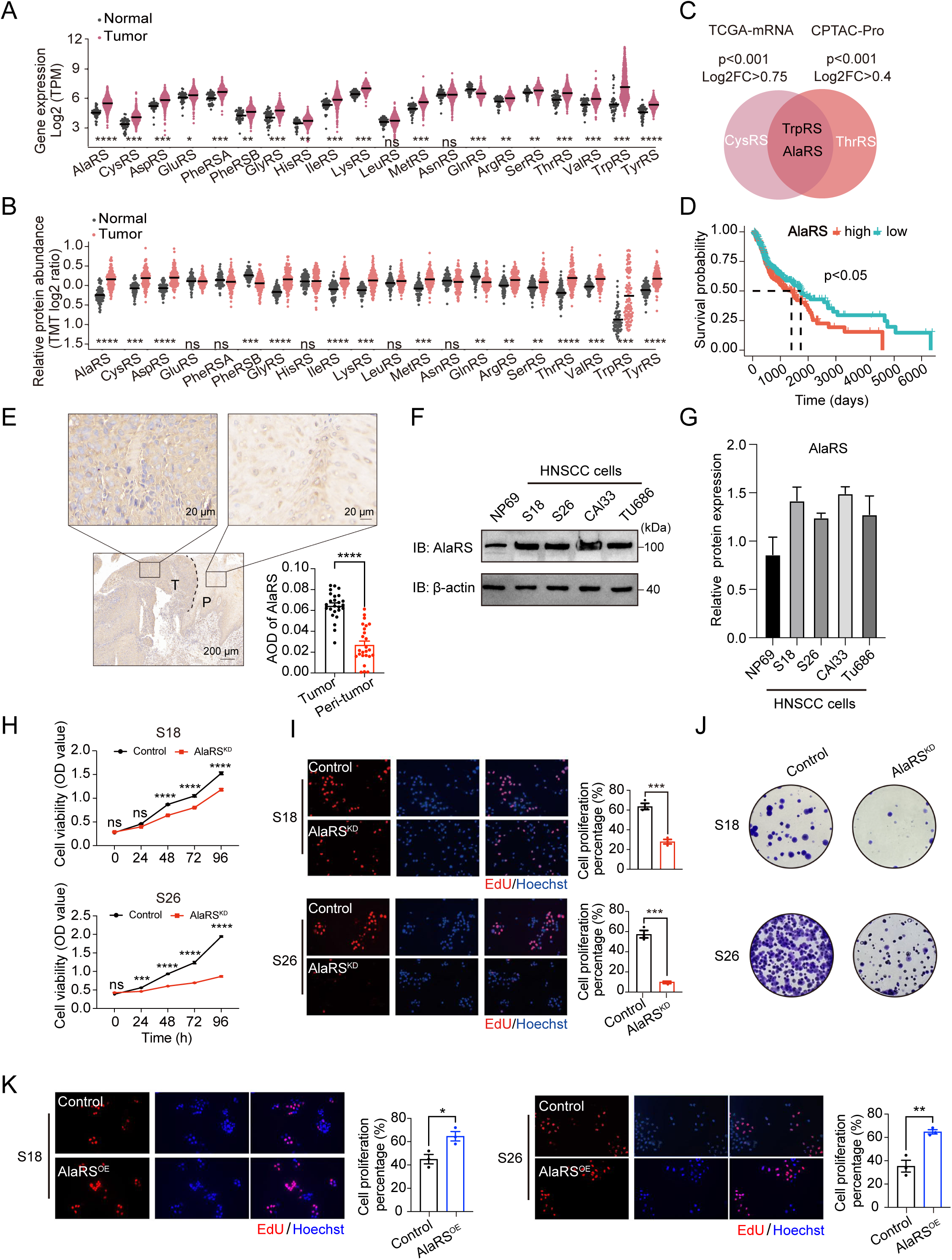
AlaRS is upregulated in HNSCC and correlates with poor prognosis and enhanced tumor cell proliferation. **A)** The mRNA expression levels of aaRSs in tumors (magenta) verus normal tissues (gray) from TCGA, analyzed by Wilcoxon rank-sum test. **B)** The protein expression levels of aaRSs in tumors (salmon) verus normal tissues (gray) from CPTAC dataset, analyzed by Wilcoxon rank-sum test. **C)** Venn diagram showing the overlap of the top three upregulated aaRSs identified from TCGA and CPTAC databases. **D)** Kaplan-Meier overall survival curve for HNSCC patients stratified by high versus low AlaRS expression (Log-rank test, p < 0.05). **E)** IHC staining of AlaRS in matched tumor and adjacent non-tumor tissues from HNSCC patients. Semi-quantitative comparison of average optical density (AOD) between the two groups (n = 25). Scale bars, 20 or 200 μm. **F-G)** Western blot analysis **(F)** and quantification **(G)** of AlaRS protein levels in various HNSCC cell lines and the normal nasopharyngeal epithelial cell line NP69 cells. **H)** Cell proliferation assessed by CCK-8 assay in S18 and S26 cells stably expressing control or AlaRS-targeting shRNA (AlaRS^KD^) over 96 h (n = 3). **I)** EdU staining of control and AlaRS^KD^ cells. Representative images and quantification of EdU-positive cells are shown (n = 3). **J)** Colony formation assay of control and AlaRS^KD^ stable cell lines after 2 weeks. Colonies were stained with crystal violet and representative images are shown. **K)** EdU staining of control and AlaRS^OE^ stable cell lines Representative images and quantification are shown (n = 3). ns, not significant, *p < 0.05, **p < 0.01, ***p < 0.001, ****p < 0.0001.

We further verified these observations in 15 HNSCC patients via immunohistochemistry (IHC) (**Table S1**). The staining demonstrated markedly higher AlaRS expression in tumor tissues relative to adjacent non-tumor tissues (**Figure 1E**). This clinical validation was reinforced by western blot analysis comparing multiple HNSCC cell lines (S18, S26, CAL33, and TU686) with the immortalized nasopharyngeal epithelial line NP69, which confirmed robust AlaRS protein accumulation in the tumorigenic cell lines (**Figure 1F, G**).

To directly assess the functional contribution of AlaRS on HNSCC growth, we generated stable AlaRS-overexpressing and knockdown cell lines (**Figure S2A-D**). Subsequent investigations using DNA synthesis and clonogenic assays revealed that AlaRS knockdown led to reduced cell viability (**Figure 1H-J**). Conversely, cells with AlaRS overexpression exhibited enhanced growth behavior (**Figure 1K; Figure S2E**). Collectively, these integrated in silico and *in vitro* data demonstrate that AlaRS acts as a key driver of tumor cell proliferation in HNSCC.

### 2.2. AlaRS modulates cell cycle related gene expression in HNSCC cells

To elucidate the molecular mechanism regulated by AlaRS, we further performed RNA sequencing (RNA-seq). Comparative transcriptomic analysis between the AlaRS knockdown (AlaRS^KD^) and control group, followed by Kyoto Encyclopedia of Genes and Genomes (KEGG) pathway enrichment analysis, revealed several altered pathways, including "Ribosome" and "Ribosome biogenesis in eukaryotes" (**Figure 2A**). Notably, the "Cell cycle" pathway also emerged as a key pathway of interest. A volcano plot revealed a significant upregulation of the cell cycle inhibitor *CDKN1A/p21* in AlaRS^KD^ group (**Figure 2B**). Further qPCR analysis verified the downregulation of several cell cycle-related genes, including *E2F5*, *CDK1*, *HDAC2*, and *CCNA2* in AlaRS^KD^ group (**Figure 2C**). Immunoblot analysis confirmed significant upregulation of p21 protein levels, along with a reduction in phosphorylated retinoblastoma tumor suppressor protein (p-Rb), underscoring their role as key regulators in modulating HNSCC cell cycle progression in response to AlaRS knockdown (**Figure 2D**).

**Figure 2.**
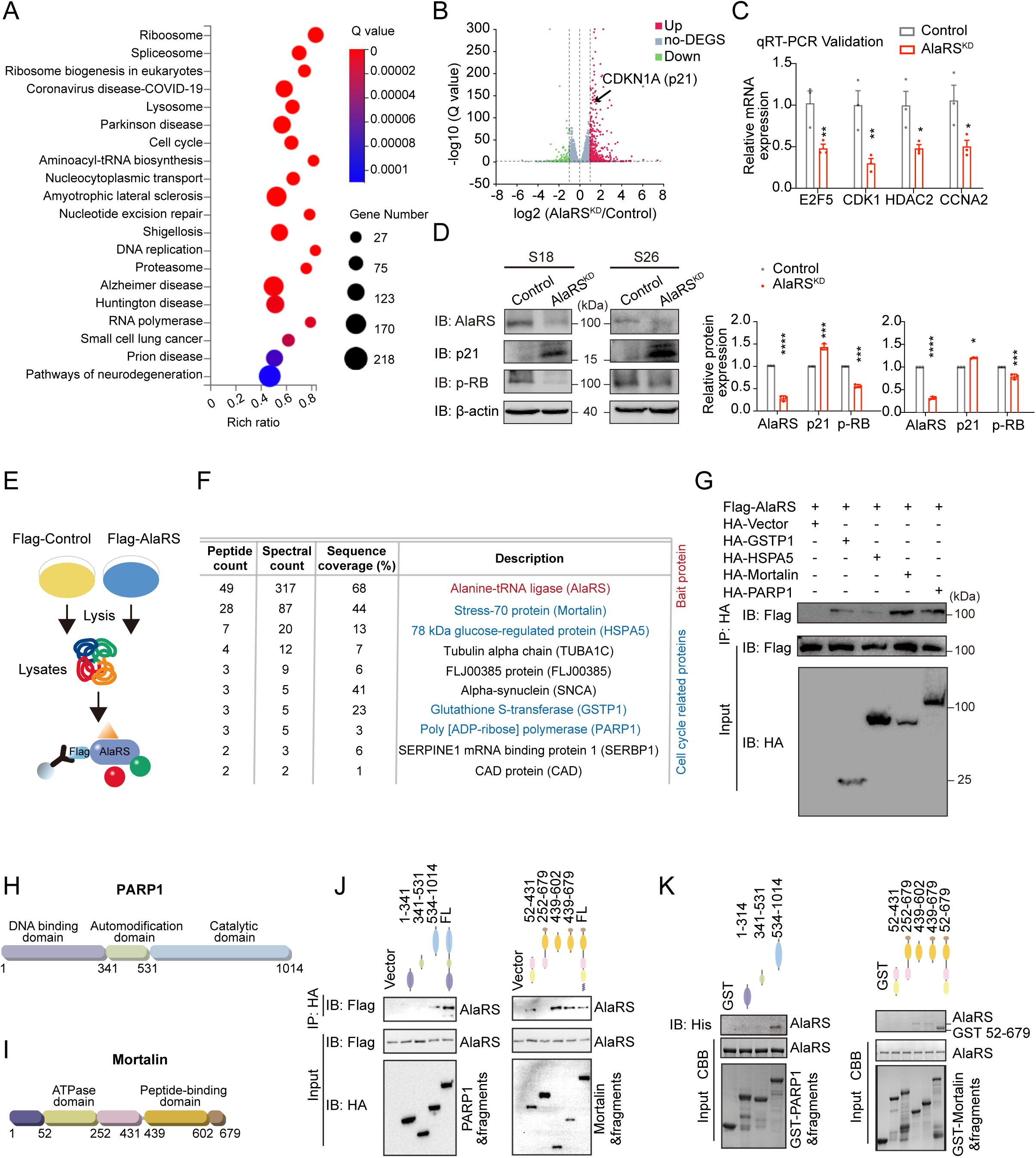
AlaRS modulates cell cycle related gene expression and interacts with PARP1 and Mortalin. **A**) Kyoto Encyclopedia of Genes and Genomes (KEGG) pathway enrichment analysis of differentially expressed genes (DEGs) following AlaRS knockdown compared to control cells. **B**) Volcano plots of DEGs (|log_2_ (fold change) | ≥ 1, FDR < 0.05) from RNA-seq of AlaRS^KD^ verus control S26 cells. The y-axis represents significance (-log_10_(Q value)), while the x-axis represents log_2_ (fold change) value. Dotted lines denote thresholds. **C**) qRT-PCR validation of selected cell cycle-related DEGs from RNA-seq (AlaRS^KD^ verus control S26 cells). Data were normalized to *ACTB* and are presented as mean ± SEM. **D**) Western blot and quantitative analysis of p21 and p-RB protein levels in S18 and S26 cells stably expressing control or AlaRS-targeting shRNA (n=3). **E**) Schematic workflow for immunoprecipitation-mass spectrometry (IP-MS) to identify AlaRS-interacting proteins from Flag-AlaRS-overexpressing HEK293T cells. **F**) Top 10 most abundant proteins specifically co-immunoprecipitated with AlaRS in HEK293T cells, ranked by peptide/spectral counts. **G**) Co-IP validation of AlaRS interaction with HA-tagged GSTP1, HSPA5, Mortalin, and PARP1 in HEK293T cells. **H-I)** Domain schematics of human PARP1 **(H)** and Mortalin **(I)**. **J-K)** Mapping of AlaRS-binding regions on PARP1 and Mortalin by Co-IP **(J)** and GST pull-down **(K)** using truncated constructs. *p < 0.05, **p < 0.01, ***p < 0.001, ****p < 0.0001.

### 2.3. AlaRS interacts with PARP1 and Mortalin in HNSCC cells

To identify AlaRS-interacting proteins, we overexpressed Flag-tagged AlaRS and performed anti-Flag immunoprecipitation followed by mass spectrometry (IP-MS) (**Figure 2E**). Proteomic analysis identified 75 candidate interactors specifically enriched with AlaRS (**Figure S3**), including GSTP1, HSPA5, Mortalin (also known as HSPA9), and PARP1, which are known to be associated with the cell cycle (**Figure 2F**). Co-IP assays confirmed that AlaRS has a stronger binding affinity with PARP1 and Mortalin, compared to others (**Figure 2G**).

Domain-mapping experiments were performed by co-expressing Flag-AlaRS with HA-tagged truncations of PARP1 or Mortalin. The catalytic domain of PARP1 (residues 533-1014) and the peptide-binding domain of Mortalin (residues 439-602) were able to pull down AlaRS (**Figure 2H-J**). Direct binding was validated using purified recombinant truncated proteins in pairwise GST pull-down assays. Consistently, the results revealed that the PARP1 fragment interacting with AlaRS corresponds to residues 533-1014, while the minimal Mortalin region binding to AlaRS spans residues 439-602 (**Figure 2K**).

### 2.4. AlaRS upregulates PARP1 and Mortalin to promote HNSCC cell growth

Given that PARP1 and Mortalin are pivotal regulators of genome stability and p53 signaling respectively, we sought to determine whether these effectors mediate the oncogenic functions of AlaRS in HNSCC^28, 29^. Consistent with the upregulation of AlaRS, both PARP1 and Mortalin protein levels were markedly elevated in HNSCC cell lines and patient tumor tissues. (**Figure 3A-D**). To determine whether their expression is regulated by AlaRS, we overexpressed AlaRS in two HNSCC cell lines and observed a corresponding increase in both PARP1 and Mortalin levels (**Figure 3E**). Conversely, AlaRS knockdown reduced their protein levels, coupled with an upregulation of p53, confirming positive regulation by AlaRS (**Figure 3F**).

**Figure 3.**
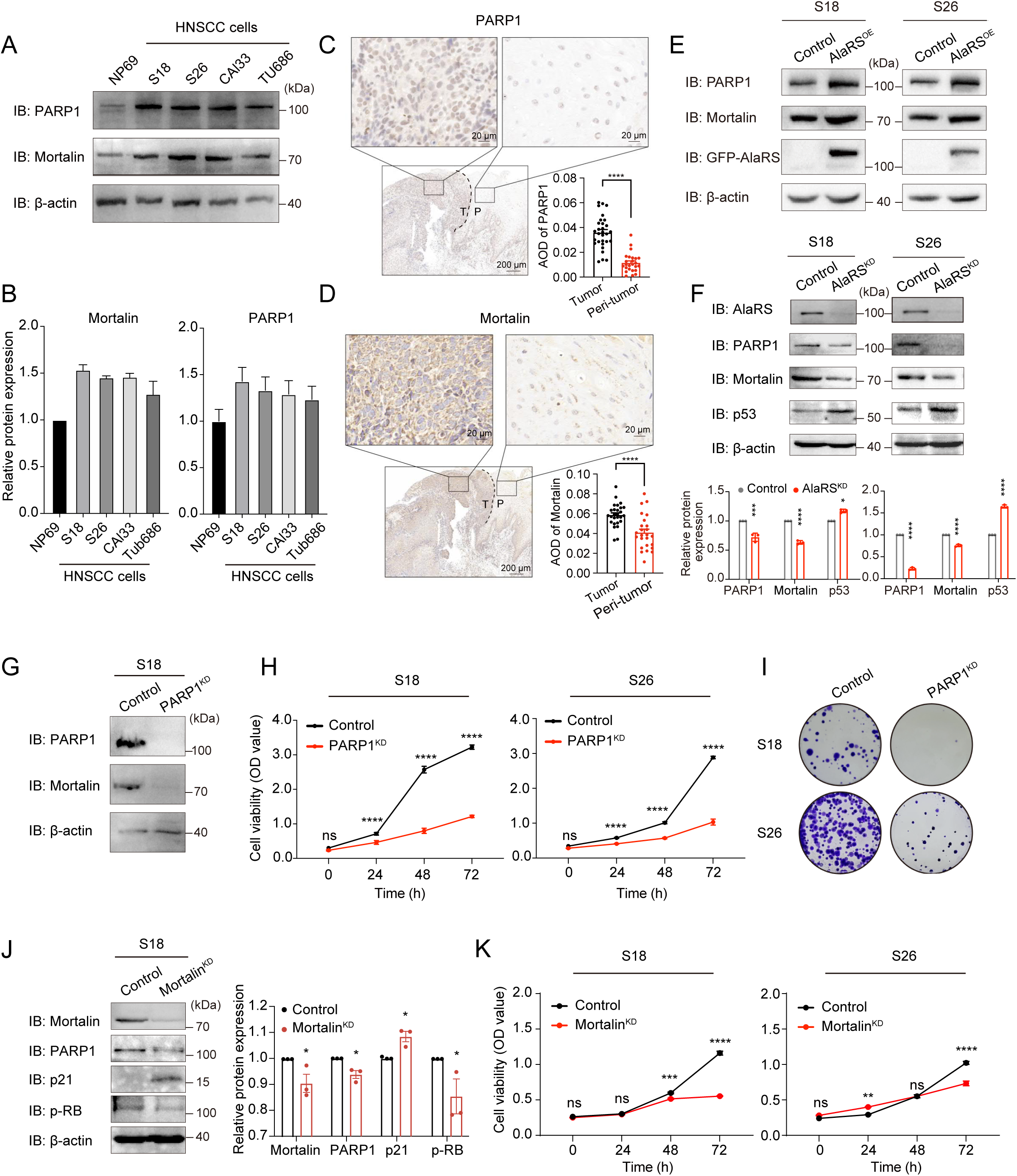
AlaRS promotes HNSCC cell growth by upregulating PARP1 and Mortalin. **A-B)** Western blot analysis **(A)** and quantification **(B)** of PARP1 and Mortalin protein levels in different HNSCC cell lines and NP69 cells. **C-D)** IHC staining of PARP1 **(C)** and Mortalin **(D)** in tumors and adjacent tissues from HNSCC patients. Semi-quantitative analysis compared AOD (n = 25). Scale bars, 20 or 200 μm. **E)** Western blot analysis of PARP1 and Mortalin protein levels following AlaRS overexpression in S18 and S26 cells. **F)** Western blot analysis and quantification of Mortalin, PARP1, and p53 protein levels following AlaRS knockdown in S18 and S26 cells (n = 3). **G)** Western blot analysis of PARP1, AlaRS, and Mortalin protein levels in PARP1^KD^ cells. **H)** Cell proliferation of PARP1^KD^ cell lines assessed by CCK-8 assay (n = 3). **I)** Representative images of colony formation assays with PARP1^KD^ cell lines. **J)** Western blot and quantification of Mortalin, PARP1, p21, and p-RB protein levels in Mortalin^KD^ cells (n = 3). **K)** Cell proliferation of Mortalin^KD^ cell lines assessed by CCK-8 assay (n = 3). ns, not significant, *p < 0.05, **p < 0.01, ***p < 0.001, ****p < 0.0001.

Co-IP and GST pull-down assays demonstrated a direct interaction between PARP1 and Mortalin (**Figure S4A, B**), consistent with prior report^30^. To determine if AlaRS functions through PARP1, we generated PARP1-knockdown cells. PARP1 depletion reduced Mortalin protein levels (**Figure 3G**) and suppressed cell proliferation or colony formation, underscoring its essential role in HNSCC growth (**Figure 3H, I**). To further delineate the role of Mortalin, we silenced Mortalin expression. Mortalin knockdown decreased PARP1 levels, elevated p21, and reduced p-RB, indicating cell cycle arrest (**Figure 3J**). Functionally, Mortalin depletion markedly impaired cell proliferation (**Figure 3K**), confirming its importance in maintaining the malignant phenotype of HNSCC cells. Collectively, these findings indicate that AlaRS facilitates HNSCC progression by coordinately upregulating PARP1 and Mortalin.

Notably, knockdown of PARP1 resulted in a more significant reduction in Mortalin protein levels compared to the decrease in PARP1 levels observed following Mortalin knockdown (**Figure 3G, J**). Functionally, PARP1 depletion also led to a stronger suppression of cell proliferation compared to Mortalin knockdown, particularly after 48 h in CCK-8 assays (**Figure 3H, K**). This asymmetric interdependence suggests that PARP1 plays a central role within this interaction, likely regulating Mortalin stability, whereas Mortalin appears to act primarily as a co-regulatory factor. Pull-down assays further demonstrated that addition of purified Mortalin did not enhance the AlaRS-PARP1 interaction (**Figure S4C**), revealing that Mortalin is a downstream effector of PARP1 stabilization rather than a structural scaffold. Consequently, we focused subsequent investigations on the direct regulation of PARP1 by AlaRS.

### 2.5. AlaRS protects PARP1 from proteasomal degradation via lactylation

To investigate how AlaRS regulates PARP1 abundance, we first examined the mRNA levels of PARP1 following AlaRS knockdown and found no significant change, suggesting that AlaRS does not affect PARP1 transcription (**Figure 4A**). We then assessed the effect of AlaRS on PARP1 protein stability using cycloheximide (CHX), an inhibitor of protein synthesis. Western blot analysis revealed that AlaRS knockdown shortened the half-life of PARP1, suggesting that the decrease in PARP1 protein levels resulted from promoted degradation through a specific proteolytic pathway (**Figure 4B**). Indeed, treatment with the proteasome inhibitor MG132 rescued PARP1 levels depleted by AlaRS knockdown, supporting a role for AlaRS in suppressing proteasome-mediated degradation of PARP1 (**Figure 4C**). Consistently, AlaRS overexpression reduced PARP1 ubiquitination. These results indicate that AlaRS stabilizes PARP1 by inhibiting its ubiquitin-dependent proteasomal degradation (**Figure 4D**).

**Figure 4.**
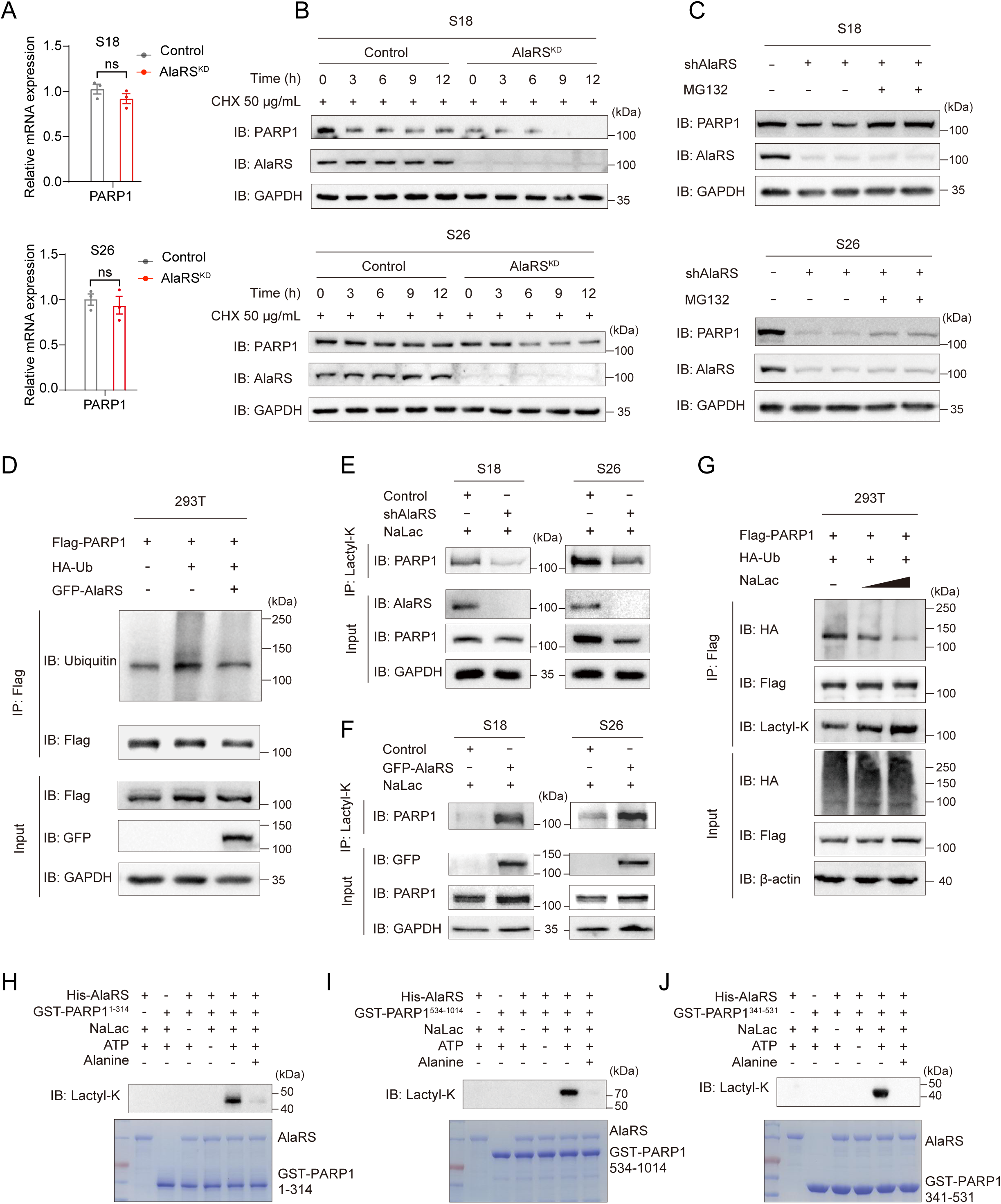
AlaRS stabilizes PARP1 by inhibiting its ubiquitin-proteasome-mediated degradation. **A)** Relative *PARP1* mRNA expression in S18 and S26 cells stably expressing control or AlaRS-targeting shRNA. ns, not significant. **B)** Half-life of PARP1 protein in S18 and S26 cells after AlaRS knockdown, determined by cycloheximide (CHX, 50 μg/mL) chase assay. **C)** Western blot analysis of PARP1 protein levels in AlaRS^KD^ S18 and S26 cells treated with MG132 (10 μM, 3 h). **D)** Co-IP analysis of PARP1 ubiquitination in HEK293T cells overexpressing AlaRS. **E-F)** Western blot analysis of whole-cell lysates (WCL) and anti-lactyl-lysine immunoprecipitates from S18 and S26 cells with AlaRS knockdown **(E)** or overexpression **(F)**, following stimulation with 24 mM lactate for 24 h. **G)** Western blot analysis of PARP1 ubiquitination in HEK293T cells co-transfected with Flag-PARP1 and HA-Ub, with or without lactate (12 or 24 mM) stimulation for 24 h. **H-J)** *In vitro* lactylation assays detecting AlaRS-induced lactylation of GST-PARP1 fragments with a pan-Klac antibody. Coomassie brilliant blue (CBB) staining shows purified His-AlaRS and GST-PARP1 fragments.

Given the reported lysine lactyltransferase activity of AlaRS^31, 32^, we hypothesized that lactylation might mediate this regulatory effect. We found that AlaRS catalyzed the PARP1 lactylation in HNSCC cells (**Figure 4E, F**). Notably, increased PARP1 lactylation was accompanied by decreased ubiquitination, suggesting that lactylation competitively inhibits ubiquitination and thereby enhances PARP1 stability (**Figure 4G**). Together, these results demonstrate that AlaRS-mediated lactylation of PARP1 prevents its degradation through the ubiquitin-proteasome pathway, thereby promoting PARP1 protein stability. To determine whether AlaRS directly catalyzes lysine lactylation of PARP1, we performed *in vitro* lactylation assays by incubating purified AlaRS with recombinant PARP1 fragments (**Figure 4H-J**). PARP1 lactylation was detected only in the presence of sodium lactate and ATP. Addition of alanine abolished this modification.

### 2.6. Identification of K249 and K667 as critical lactylation sites on PARP1

To identify specific modification sites within PARP1, we employed LC-MS/MS and revealed 22 potential lactylation sites (**Table S2**). For functional validation, we successfully purified 10 site-directed PARP1 mutants, which covered 15 of the putative lactylation sites (7 sites excluded due to protein instability). In subsequent *in vitro* lactylation assays, mutations at the K249 and K667 residues in the truncated PARP1 protein resulted in the most pronounced alterations in lactylation levels (**Figure 5A, B**). To validate the functional importance of these two residues in the full-length protein within cells, we generated and analyzed individual (K249R or K667R) and combined (K249R/K667R) mutants of PARP1. The single mutants did not exhibit significant changes in lactylation, while the dual mutation showed a pronounced reduction in lactylation (**Figure 5C**).

**Figure 5.**
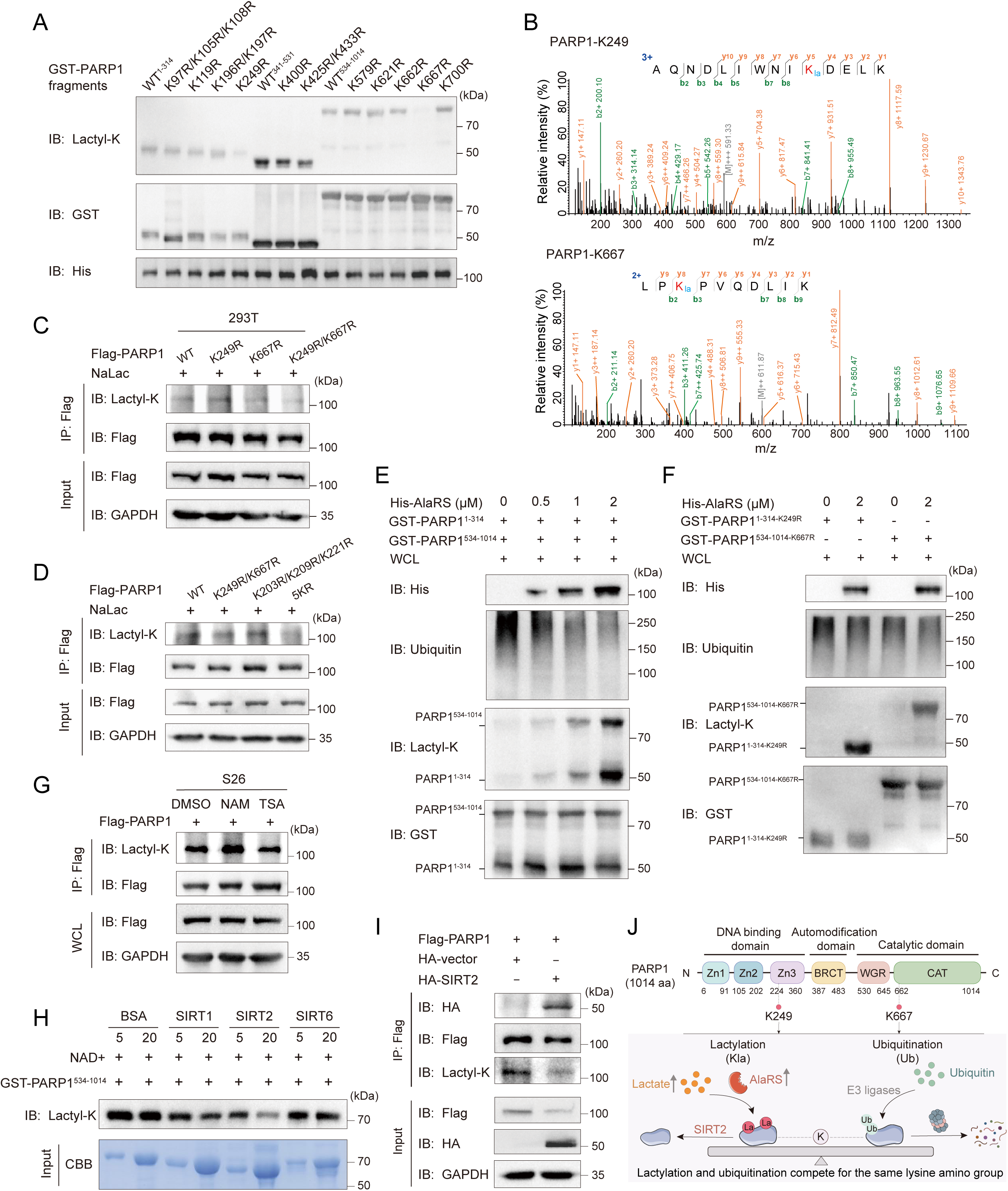
AlaRS-mediated lactylation of PARP1 antagonizes ubiquitination at K249 and K667. **A)** *In vitro* lactylation assays using purified His-AlaRS and truncated GST-PARP1 proteins with site-directed mutations at putative lysine residues. **B)** Mass spectrometry identification of *in vitro* lactylated peptides from PARP1 fragments containing K249 or K667. **C-D)** Western blot analysis of PARP1 lactylation in HEK293T cells transfected with the indicated PARP1 mutants. 5KR: K203R/K209R/K221R/K249R/K667R. **E-F)** Competitive lactylation and ubiquitination assays using purified wild-type **(E)** or mutant **(F)** PARP1 fragments incubated with increasing concentrations of AlaRS protein, ATP (5 mM) and sodium lactate (5 mM) for 2 h at 25℃, followed by exposure to a cell lysate-based ubiquitination system. **G)** Western blot analysis of Flag-PARP1 lactylation levels in S26 cells treated with NAM or TSA. **H)** *In vitro* delactylation assay using recombinant sirtuin proteins and lactylated PARP1 in the presence of NAD+ for 3 h at 37℃. **I)** Western blot analysis of Flag-PARP1 lactylation in S26 cells overexpressing SIRT2. **J)** Schematic model mapping K249 and K667 within the PARP1 domain architecture and depicting the competitive interplay between lactylation and ubiquitination at these specific residues.

The key lactylation residues K249 and K667 on PARP1 intersect with known ubiquitination sites^33–35^, implicating them in a competitive crosstalk that may dynamically regulate protein stability (**Figure S5A**). We next functionally assessed other residues subject to both modifications. Notably, a triple mutant (K203R/K209R/K221R)^35, 36^ exerted a markedly less significant effect on lactylation than K249R/K667R double mutant, underscoring that K249 and K667 are the dominant lactylation sites essential for regulating PARP1 stability (**Figure 5D; Figure S5B**).

Subsequently, to explore whether K249 and K667 are hubs for competitive lactylation-ubiquitination crosstalk, we first incubated purified PARP1 fragments with increasing concentrations of AlaRS to promote site-specific lactylation. This pre-lactylated system was then subjected to a ubiquitination reaction using cell lysates containing the necessary enzymes and substrates. Analysis of the GST-enriched PARP1 fragments revealed that higher concentrations of AlaRS correlated with increased PARP1 lactylation and a concomitant decrease in its ubiquitination, suggesting competitive occupancy of shared lysine residues (**Figure 5E**). Strikingly, when K249 or K667 was mutated, the addition of AlaRS induced only minimal lactylation (attributable to other minor sites) and failed to substantially suppress ubiquitination. This highlights that AlaRS primarily regulates the lactylation-ubiquitination balance on PARP1 through the specific sites K249 and K667 (**Figure 5F**).

We next sought the delactylase that counteracts AlaRS. In S26 cells, the pan-sirtuin inhibitor nicotinamide (NAM), but not the pan-HDAC inhibitor Trichostatin A (TSA), elevated PARP1 lactylation (**Figure 5G**), implicating sirtuins. Given that SIRT1, SIRT2, and SIRT6 are known to interact with PARP1^37–40^, we evaluated their enzymatic activities via *in vitro* delactylation assays. Among these candidates, SIRT2 exhibited the most robust NAD+ dependent removal of lactyl groups from PARP1 (**Figure 5H**). This was confirmed *in cellulo*, as SIRT2 overexpression markedly abolished PARP1 lactylation (**Figure 5I**). Collectively, these results establish SIRT2 as the primary eraser of AlaRS-mediated PARP1 lactylation. Based on these findings, we propose a mechanistic model in which AlaRS-mediated lactylation at K249 and K667, reversibly regulated by SIRT2, competitively antagonizes ubiquitination at the same residues, thereby protecting PARP1 from proteasomal degradation (**Figure 5J**).

### 2.7. CHE selectively targets AlaRS to inhibit PARP1 lactylation

A multi-step screening pipeline was employed to identify small molecules binding AlaRS and inhibiting its lactyltransferase activity (**Figure 6A**). Initially, recombinant AlaRS protein was purified and utilized in a fluorescence-based TSA^41^. We screened 5,090 small molecules, including 2,734 bioactive compounds and 2,356 FDA-approved pharmaceuticals, and identified 14 hits with Tm shifts >3.0°C (**Figure 6B**). Subsequent BLI analysis determined the binding kinetics and affinities of these candidates, identifying 8 compounds with strong binding to AlaRS (**Figure 6C; Table S3**).

**Figure 6.**
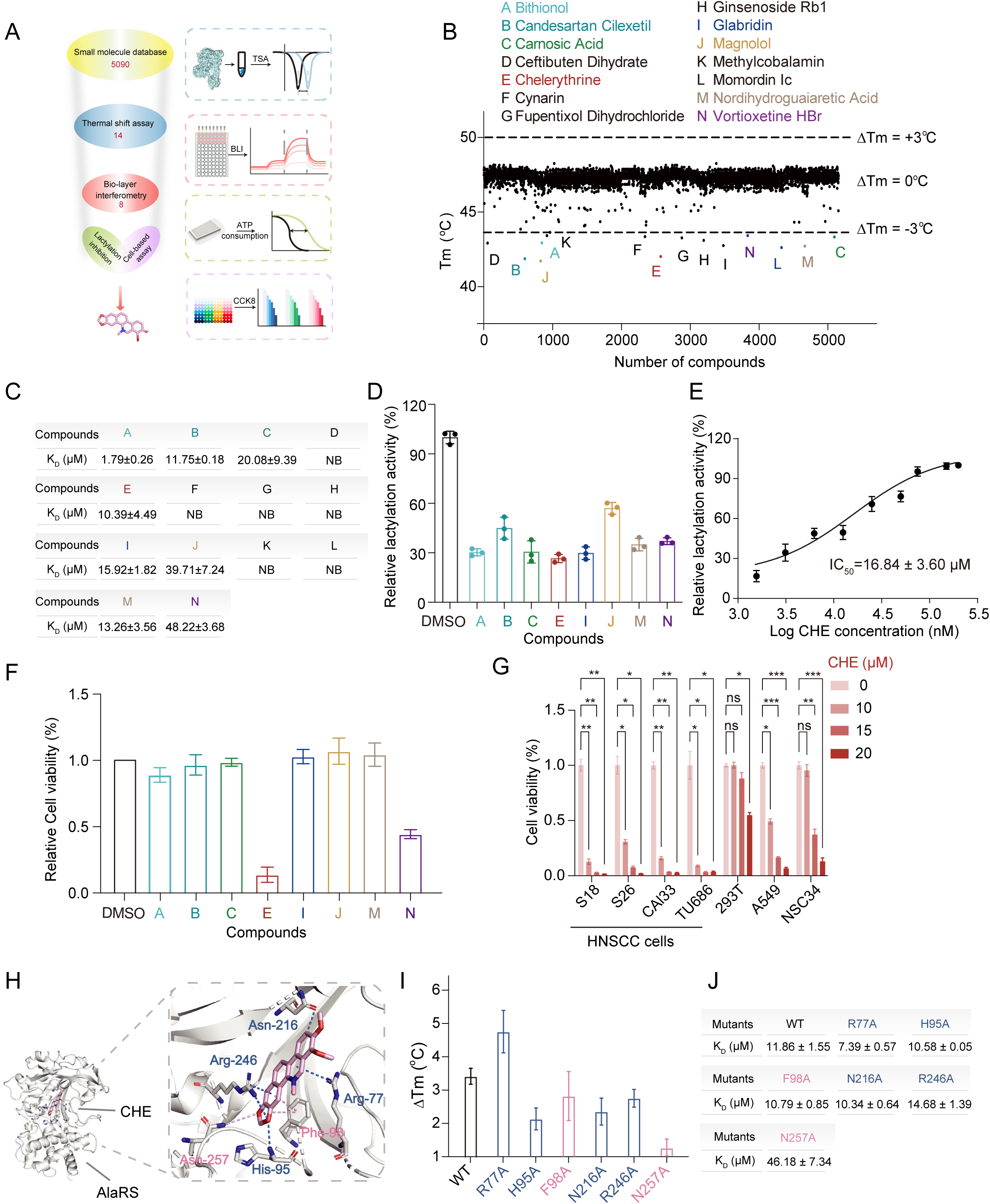
Identification of small-molecule inhibitors targeting AlaRS in HNSCC. **A)** Schematic of the multi-step screening strategy, including thermal shift assay (TSA), bio-layer interferometry (BLI), *in vitro* lactylation inhibition assay, and CCK-8 cell viability assay. **B)** TSA results for 5,090 compounds plotted by melting temperature (Tm) shift. Dots outside the gray dashed lines represent positive results. **C)** Binding affinity of selected compounds for AlaRS as measured by BLI (n = 3). Data are presented as mean ± standard deviation (SD). NB, no binding. **D)** ATP consumption assay measuring the inhibiton of AlaRS-mediated lactylation by eight candidate compounds (n = 3). **E)** Dose-dependent inhibition of AlaRS-mediated lactylation by CHE, measured by ATP consumption (n = 3). **F)** Cell viability of S18 cells treated with 10 µM candidate compounds, assessed by CCK-8 assay (n = 3). **G)** Anti-proliferative activity of CHE in HNSCC and other cell lines, evaluated by CCK-8 assay (n =3). ns, not significant, *p < 0.05, **p < 0.01, ***p < 0.001. **H)** Molecular docking model of CHE (colored sticks) bound to human AlaRS (PDB: 4XEM). **I)** TSA analysis of melting temperature shifts for six AlaRS point mutants upon CHE binding. **J)** Binding affinity of CHE to six AlaRS mutants measured by BLI (n = 3). Data are presented as mean ± SD.

To assess functional inhibition, we conducted *in vitro* lactylation assays based on ATP consumption. CHE emerged as the most potent lead, inhibiting AlaRS-mediated lactylation with an IC_50_ of 16.84 μM and effectively suppressing the site-specific lactylation of PARP1 (**Figure 6D, E; Figure S6A**). Crucially, CHE exhibited superior anti-proliferative activity specifically in HNSCC cells compared to other candidates (**Figure 6F; Figure S6B; Table S4**). To evaluate its selectivity, we measured the IC_50_ of CHE across a panel of cell lines. The IC_50_ values in four HNSCC cell lines ranged from 4.77 to 5.11 µM, whereas values in non-HNSCC lines (NSC34, A549, HEK293T) were markedly higher (17.43-21.54 µM) (**Figure 6G; Figure S6C**). This pronounced difference in sensitivity supports a selective inhibitory effect of CHE against HNSCC.

To deepen our understanding of the mechanism of CHE acting on AlaRS, we performed molecular dynamics and molecular docking. Through monitoring the RMSD and RMSF changes of CHE during the simulation, we observed that CHE was stable at the AlaRS binding site (**Figure S7A**). Docking analysis using AutoDock Vina revealed that CHE was located in the aminoacylation region (**Figure 6H**).

Structural superposition with the Ala-AMP analog, AlaSA, revealed that CHE occupies a site partially overlapping with the canonical reaction intermediate (**Figure S7B**). While several hydrogen-bonding residues are shared between CHE and AlaSA, mutational analysis pinpointed Asn257 as a critical determinant for CHE binding. Specifically, the N257A mutation abolished the CHE-induced Tm shift and increased the K_D_ value by over four-fold, whereas other conserved residue substitutions had minimal impact (**Figure 6I, J; Figure S7C**).

Interestingly, aaRSs occasionally misincorporate non-cognate amino acids due to structural similarity. For instance, AlaRS is known to mischarge serine or glycine in addition to its canonical substrate alanine, with significant functional implications^42–44^. This flexibility extends to lactate, which structurally resembles alanine and can bind within the catalytic site of AlaRS^31^. Our molecular docking revealed that the inhibitor CHE also localizes to the aminoacylation domain of AlaRS (**Figure 6H**). We further confirmed that CHE inhibits the aminoacylation activity of AlaRS in a dose-dependent manner (**Figure S7D**). Collectively, these data establish CHE as a selective, dual-function inhibitor that targets the AlaRS catalytic center to suppress HNSCC progression.

### 2.8. CHE induces PARP1 and Mortalin degradation and suppresses tumor growth *in vivo*

To validate the cellular efficacy of CHE, we examined its effect on cell viability across four HNSCC cell lines. Consistently, EdU assays, Annexin V-FITC staining, and colony formation assays indicated that CHE significantly inhibited HNSCC cell proliferation, and induced apoptosis (**Figure S8A-C**). To determine whether these effects are specifically mediated by AlaRS, we modulated AlaRS expression in S18 and S26 cells. Notably, AlaRS overexpression enhanced sensitivity to CHE (**Figure 7A**), indicating that CHE’s anti-proliferative activity is largely dependent on AlaRS. In addition, western blot analysis showed CHE treatment led to a concomitant downregulation of PARP1 and Mortalin (**Figure 7B, C**).

**Figure 7.**
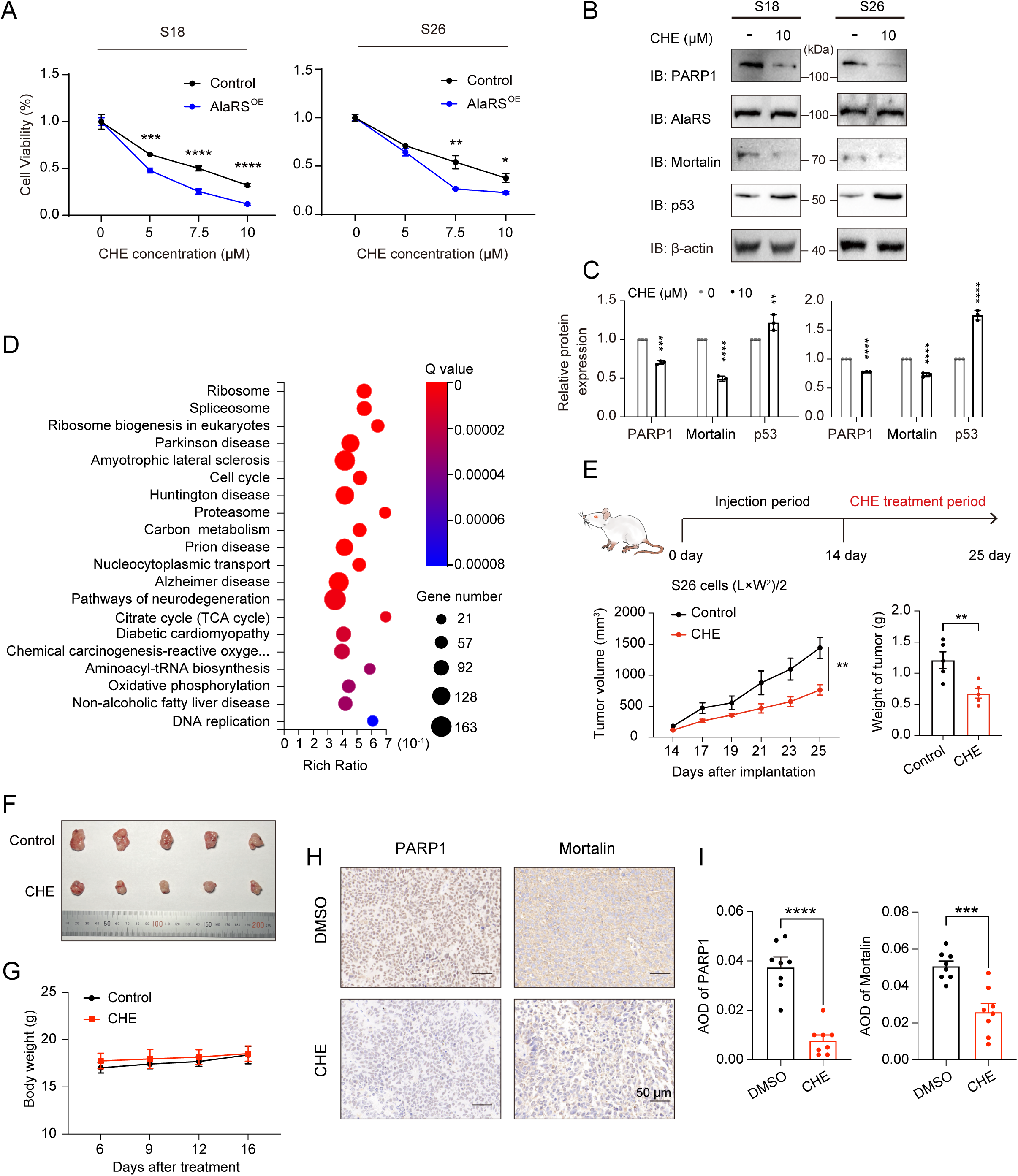
CHE treatment inhibits HNSCC cell viability and tumor growth in mice. **A)** Cell viability analysis in S18 and S26 cells stably expressing control or AlaRS^OE^ following treatment with CHE (n = 3). **B-C)** Western blot analysis and quantification of PARP1, AlaRS, Mortalin, and p53 protein levels in HNSCC cells treated with CHE (n = 3). **D)** KEGG pathway enrichment analysis of DEGs identified by RNA-seq after CHE treatment. **E-F)** Tumor volume and tumor weight **(E)** of S26 xenografts in nude mice treated with CHE (n = 5). Representative tumor images at endpoint are shown **(F)**. Data are presented as mean ± SEM. **G)** Body weight monitoring from S26 xenograft mice treated with CHE or vehicle control (n = 5). **H-I)** IHC staining **(H)** and quantification **(I)** of PARP1 and Mortalin in xenograft tumor tissues (n = 8). Scale bars, 50 μm. *p < 0.05, **p < 0.01, ***p < 0.001, ****p < 0.0001.

RNA sequencing of CHE-treated S26 cells followed by KEGG pathway enrichment analysis identified significant enrichment in “Ribosome”, “Ribosome biogenesis in eukaryotes”, and “Cell cycle” pathways (**Figure 7D**). These findings support that CHE acts on AlaRS to reduce the lactylation of PARP1, thereby decreasing stability of PARP1 and Mortalin and consequently impacting cell cycle progression.

The *in vivo* antitumor efficacy of CHE was assessed using a subcutaneous xenograft model established with S26 cells. Over 11 days of treatment, CHE significantly suppressed tumor growth and reduced endpoint tumor weight compared to vehicle control (**Figure 7E, F**). Importantly, no significant body weight loss was observed in CHE-treated mice (**Figure 7G**), suggesting that at the therapeutic dose, partial inhibition of AlaRS’s canonical aminoacylation activity does not compromise normal tissue homeostasis, as residual tRNA charging remains sufficient. Western blotting and IHC of harvested tumors confirmed that CHE downregulated PARP1 and Mortalin protein levels *in vivo* (**Figure 7H, I; Figure S9**). Together with our *in vitro* findings, these results demonstrate that CHE is a potent and well-tolerated antitumor agent in HNSCC that acts primarily through direct targeting of AlaRS.

## 3. Discussion

Lactate is widely recognized as a glycolytic metabolite that supports energy production. In cancer, the Warburg effect drives increased intracellular lactate accumulation and acidification of the tumor microenvironment. Beyond its metabolic role, lactate also functions as a signaling molecule, influencing various biological processes such as immune regulation, angiogenesis, fibrosis, and tumor proliferation^45–47^. A key mechanism through which lactate conveys these signals is protein lactylation, a post-translational modification that uses lactate as a substrate. This modification represents an important regulatory pathway in early tumor progression. Our study identifies AlaRS as a pivotal lactyltransferase that translates metabolic flux into oncogenic signals. By demonstrating that AlaRS-mediated lactylation sustains HNSCC progression, we provide a mechanistic foundation for understanding how glycolytic hyperactivity is directly coupled to protein stability and tumor growth.

AaRSs are increasingly recognized as multifaceted contributors to tumor progression. Beyond their essential role in translation, certain aaRSs perform non-canonical functions, such as nutrient-sensing through lysine aminoacylation of target proteins, thereby regulating cellular processes and contributing to tumorigenesis^48, 49^. These diverse roles position aaRSs as promising therapeutic targets in cancer^21^. In particular, AlaRS is significantly overexpressed in multiple malignancies, including gastric^32^ and duodenal cancers^50^. Our findings expand the regulatory repertoire of aaRSs by characterizing AlaRS as a “writer” of lactylation. This dual-functional capacity, coupling tRNA aminoacylation with protein lactylation, creates a metabolic switch. While AlaRS maintains a high affinity for alanine to ensure translation under physiological conditions, the profound accumulation of lactate in the tumor microenvironment (typically 10-40 mM) reaches the kinetic threshold required for Lac-AMP formation^51, 52^. This high substrate availability drives AlaRS to prioritize its lactyltransferase activity, effectively coupling protein stability to the cell’s metabolic status and ensuring oncoprotein accumulation under metabolic stress.

We identify PARP1, a cornerstone of DNA repair and genome integrity, as a primary substrate for AlaRS in HNSCC. Although PARP1 is a well-validated clinical target, its regulation via lactylation was previously unknown. The discovery of this AlaRS-PARP1 regulatory axis suggests a metabolic escape mechanism where tumor cells stabilize PARP1 to bypass p53-mediated growth arrest. Furthermore, the direct interaction between PARP1 and the chaperone Mortalin creates a stabilizing complex, providing a rational explanation for the concomitant elevation of these proteins in HNSCC. This axis represents a potential therapeutic vulnerability that could be exploited to sensitize tumors to DNA-damaging agents or metabolic inhibitors.

A critical revelation of our study is the competitive interplay between lactylation and ubiquitination at residues K249 and K667 of PARP1. This PTM crosstalk represents a sophisticated level of protein fate determination. Interestingly, the biological outcome of AlaRS-mediated lactylation appears to be context- and substrate-dependent. While AlaRS stabilizes YAP in gastric cancer and PARP1 in HNSCC, it promotes the degradation of YTHDC1 in bladder cancer^32, 53^. This substrate-specific divergence suggests that AlaRS-mediated lactylation can either mask or recruit ubiquitin ligases depending on the local structural environment of the target lysine. In HNSCC, the “lactyl-shield” on PARP1 effectively blocks proteasomal degradation, locking the cell in a proliferative state.

A critical concern for targeting AlaRS is the potential for systemic toxicity, given its essential role in canonical tRNA charging. However, our findings reveal a viable therapeutic window for CHE driven by differential cellular dependency. Highly glycolytic HNSCC cells depend on AlaRS-mediated lactylation to stabilize key oncoproteins like PARP1 in a lactate-rich microenvironment. In contrast, normal tissues maintain low baseline lactate levels and rely on AlaRS primarily for normal protein synthesis. This biological divergence translates into a distinct drug sensitivity. Our observation that normal cell lines are significantly more resistant to CHE (**Figure 6G**) and that CHE-treated mice exhibit no overt organ toxicity or weight loss (**Figure 7F**) supports the feasibility of targeting AlaRS. Thus, CHE exploits a context-dependent tumor vulnerability, fatally disrupting the cancer-specific lactylation pathway while sparing sufficient tRNA charging for normal tissue homeostasis.

Although CHE serves as a valuable compound by concurrently inhibiting both lactyltransferase and canonical aminoacylation activities, its broader clinical application requires further refinement. Our structural data identifying N257 as a key residue for CHE binding provides a blueprint for medicinal chemistry efforts. Given that the catalytic pocket for lactyl-AMP and alanyl-AMP formation is structurally conserved, achieving biochemical selectivity at the active site remains challenging. Future strategies should therefore focus on targeting the specific protein-protein interaction interfaces between AlaRS and its protein substrates (e.g., PARP1), which likely differ from the tRNA-binding surfaces. Such non-canonical-specific inhibitors could selectively disrupt the tumor-promoting lactylation axis while preserving essential translation, further widening the therapeutic window.

In summary, our findings reveal that AlaRS links lactate accumulation to PARP1 stabilization through site-specific lactylation at K249 and K667, thereby driving HNSCC progression via the p53/p21 axis. Pharmacological inhibition of AlaRS lactyltransferase activity by CHE reduces PARP1 levels and suppresses tumor growth in preclinical models (**Figure 8**), nominating the AlaRS-PARP1 lactylation axis as a candidate therapeutic target in HNSCC.

**Figure 8.**
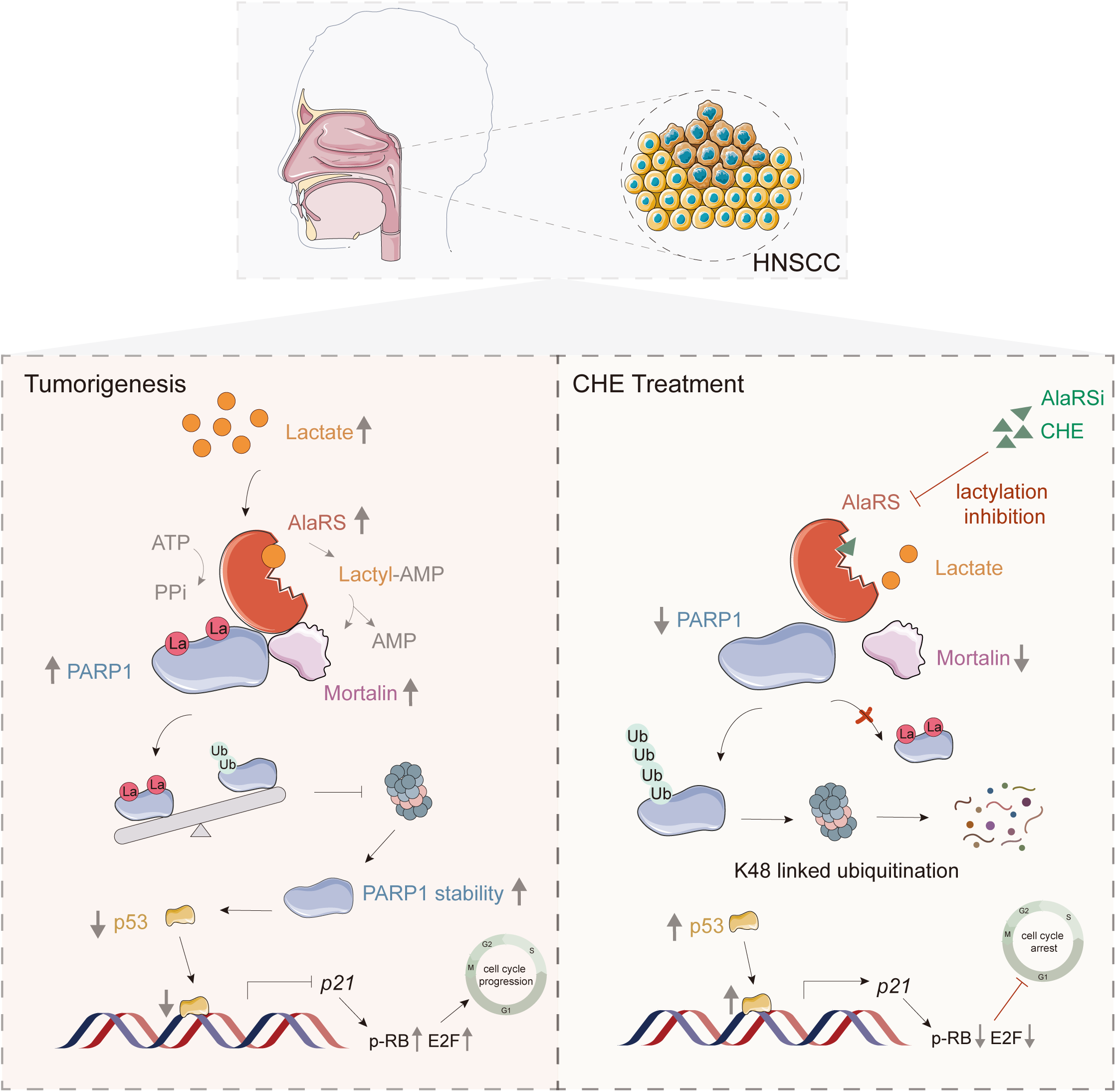
Schematic model of AlaRS-mediated PARP1 lactylation in HNSCC progression. AlaRS is upregulated in HNSCC and acts as a lactyltransferase, catalyzing site-specific lactylation of PARP1. This modification competitively inhibits PARP1 ubiquitination, thereby stabilizing PARP1 and promoting tumor progression via the p53/p21 axis. The compound CHE inhibits AlaRS lactyltransferase activity, suppressing HNSCC growth *in vitro* and *in vivo*.

## 4. Experimental Section Data collection and analysis

Data from a cohort of 542 patients (498 tumor and 44 normal samples) were analyzed, with RNA-sequencing expression quantified in transcripts per million (tpm). Somatic mutation and copy number variation (CNV) data were obtained from The Cancer Genome Atlas (TCGA-HNSCC) database. Protein expression data for 20 cytoplasmic aaRSs were retrieved from the PDC platform (https://pdc.cancer.gov). Differential expression analysis of both mRNAs and proteins, as well as patient survival data, was performed using R 4.2.2 software.

### Construction of shRNA expression vectors

The shRNA oligos were ordered from Sangong Biotech (Shanghai, China), and the sequence of these oligos were showed in Table S5. The system of the annealing oligos was followed by mixing both 20 μM forward and reverse oligo 5 μL, 5 μL 10 × NEB buffer 2 and ddH_2_O together (total volume is 20 μL). The program was 95°C for 4 min, 70°C for 10 min by PCR machine, and then slowly cooled down to room temperature for at least 2 hours. The recombinant plasmids (pLKO.1-shAlaRS, pLKO.1-shMortalin, pLKO.1-shPARP1 and a negative control pLKO.1-shScramble) were generated by digesting the vector with AgeI and EcoRI, followed by ligation with annealed oligos of indicated genes using T4 DNA ligase at 16°C. A 10 μL aliquot of the ligation mixture was transformed into 100 μL of DH5α competent cells. Positive clones harboring the correct plasmids were identified by sequencing.

### Lentivirus assembly and infection

High titers of lentivirus were generated by co-transfection of HEK293T cells with the viral constructs (10 µg) and the packaging plasmids (7.5 µg pMD2.G and 7.5 µg psPAX2) using linear polyethyleneimine (PEI; Lifescience). Meanwhile, shScramble lentivirus was used as a control. After transfection for 48 hours, the virus-containing supernatants was harvested by centrifugation (1,000×g, 5 min) and then filtered through a 0.45 µm syringe filter. The S18 and S26 cells were infected with the supernatants at a multiplicity of infection of 50, and polybrene was added to a final concentration of 8 µg/mL. Stably transduced cell lines were established by puromycin (5 μg/mL) selection. Discrete colonies resistant to puromycin appeared within a week. The cells were collected for mRNA or protein quantification to confirm knockdown.

### Protein expression and purification

Human AlaRS, Mortalin and their truncations were cloned into the pET-21a vector, while PARP1 and its truncations were cloned into the pGEx-6p-1 vector simultaneously. The constructs were then transformed into BL21(DE3) cells. Overnight cultures were grown in LB medium supplemented with 100 µg/mL ampicillin until saturation. The overnight culture was subsequently diluted 1/100 in LB medium and incubated at 37°C. At an optical density of OD600 of 0.4, isopropyl β-D-thiogalactopyranoside (IPTG) was added to a final concentration of 0.3 mM, and the cells were grown at 16°C for 20 hours to induce protein expression. After induction, the cells were harvested by centrifugation at 4°C, 4,000 rpm for 20 min. The resulting pellet was resuspended in lysis buffer (20 mM Tris-HCl, 300 mM NaCl, 5 mM imidazole, 1 mM phenyl-methyl-sulphonyl-fluoride (PMSF), pH 8.0). Cell lysis was achieved using an ultrasonic cell crusher, and the lysate was clarified by centrifugation at 4°C, 20,000 rpm for 1 hour. The proteins were purified using Ni-NTA beads (Cytiva), followed by chromatography on a HiTrap Q HP column (Cytiva), and finally, a HiLoad 16/60 Superdex 200 prep grade column (Cytiva). All purification steps were carried out at 4°C or on ice to maintain protein stability and activity.

### Thermal shift assay

Thermal shift assays were conducted using a StepOnePlus 7 Flex Real-Time Cycler (Applied Biosystems). The Protein Thermal Shift™ Kit dye from Thermo Fisher Scientific was employed to monitor the thermal stability of the protein by binding to its exposed hydrophobic regions. In each well of a 96-well Optical Reaction Plate (Applied Biosystems), a solution containing 5 μL of protein thermal shift buffer, 2.5 μL of diluted thermal shift dye (8×), and 12.5 μL of protein at 1 mg/mL was added. The 96-well polymerase chain reaction plates were then placed in the Life Technologies system and incubated at 25°C for 10 min. Subsequently, the samples were gradually heated from 25°C to 95°C at a rate of 1°C/min. During the thermal denaturation of the protein, the fluorescence signal of SYPRO orange at 490/530 nm excitation/emission wavelengths was recorded by the instrument at 30-second intervals. The fluorescence signal was continuously monitored and plotted against the temperature, with the midpoint of the protein unfolding transition defined as the melting temperature (Tm). To ensure accuracy, triplicate experiments were performed. The ΔTm represents the shift in Tm value between the AlaRS treated with compounds and the blank control.

### Bio-layer interferometry assay

An Octet R8 system (ForteBio, CA, USA) was used for conducting the binding experiments on the AlaRS proteins with molecules. The purified His-tagged AlaRS^1–84^ proteins (500 μg/mL) were captured via Ni-NTA biosensors (ForteBio), resulting in a saturation response of 4-5 nm after 300 s. Subsequently, the loaded biosensors were transferred into Octet buffer (PBS with 0.01% Tween 20) for 180 s to remove loose nonspecifically bound proteins and to establish a stable baseline. Then, they were immersed in diluted molecules (0.625-100 μM in assay buffer) for 120 s to provide the association signal, followed by transfer into Octet buffer to test for a disassociation signal for 100 s. Reference wells that utilized buffer instead of tested compounds were also included to correct the baseline shift. A parallel set of Ni-NTA sensors immersed only in buffer was prepared as the negative reference controls to correct the non-specific binding of the compounds to the biosensor surface. The signals were analyzed by a double reference subtraction protocol to deduce nonspecific and background signals and signal drifts caused by biosensor variability. All assays were performed by a standard protocol in 96-well black plates with a total volume of 200 μL/well at 30°C. The generated data were analyzed using the ForteBio Data Analysis software for correction and curve fitting.

### Lactylation activity inhibition assay

The ATP consumption assay was utilized to assess the inhibitory effect of the compounds on the lactylation activity of AlaRS. The 30 μL reaction mixture consisted of 1 μM AlaRS, 10 μM ATP, 5 mM sodium lactate, 50 mM HEPES pH 7.5, 150 mM NaCl, 50 mM KCl, 25 mM MgCl_2_, and 0.1% bovine serum albumin (BSA), along with 40 μM various compounds. The reaction mixture was incubated at room temperature for 15 min. After incubation, 10 μL of the reaction reagent was transferred to a 384-well microplate, and lactylation activity of AlaRS was detected by adding 10 μL of the ATP Assay Kit (#S0026, Beyotime, Shanghai, China). Luminescence measurements were performed using a Synergy HTX multimode microplate Reader (BioTek Inc, USA). The inhibitory rate was determined based on three independent assays, and the average value was calculated to provide a reliable assessment of the inhibitory activity of each compound.

### Cell lines and cell culture

Immortalized Human Nasopharyngeal Epithelial Cells (NP69SV40T) purchased from Meisen were cultured in complete medium. S18 and S26 cell lines were cultured in RPMI 1640 medium (Corning, New York, NY, USA) supplemented with 1% penicillin/streptomycin and 5% fetal bovine serum (FBS, Gibco, Carlsbad, CA, USA). CAl33, TU686, HEK293T, C2C12, A549 and NSC34 cells were cultured in Dulbecco’s Modified Eagle Medium (DMEM, Gibco, Carlsbad, CA, USA) supplemented with 1% penicillin/streptomycin and 10% FBS. All cell lines were validated by short-tandem-repeat analysis and were routinely tested for *Mycoplasma* contamination. All cell lines used in this study were incubated at 37°C in a humidified incubator containing 5% CO_2_.

### Cell viability measurement (CCK-8 Assay)

The Cell Counting Kit-8 (CCK-8) assay was used to test cell viability according to the manufacturer’s protocol. In brief, cells were plated at an initial density of 8×10^3^ cells/well in a 96-well plate overnight and treated with various concentrations of different molecules the next day. After culturing for 24 hours, 10 μL of CCK-8 reagent (GlpBio, Montclair, CA, USA) was added and incubated at 37°C and 5% CO_2_ for 2 hours. The absorbance was measured at 450 nm with a multimode microplate reader.

### EdU labeling and staining

Cells were seeded on slides and cultured at 37°C and 5% CO_2_. After molecules treatment, EdU (final concentration:10 μM; #C0075, Byeotime, Shanghai, China) was added to the medium and incubated for 2 hours. The cells were fixed with 4% paraformaldehyde (PFA) for 15 min at room temperature, then sealed and permeabilized with blocking solution (BS) (0.5% BSA, 0.1% Triton X-100). 100 μL of Click Additive Solution was then added to the sample for further incubation for 30 min in a dark and humidified chamber at 37°C. After three times PBS rinse, Hoechst was used to stain the nuclei. A mounting medium (Sigma, St. Louis, MO, USA) was used to seal the plates. Finally, the images were observed under a fluorescence microscope (ZEISS, Oberkochen, Germany).

### Colony formation assay

Cells were seeded at 200 cells/well to the bottom of 6-well plates. After attachment, cells were incubated with the different concentrations of CHE for 24 hours. The medium was placed with fresh medium and cells were cultured for another 14 days until the visible colonies were observed. The colonies were fixed with 4% PFA and stained with crystal violet staining solution (Byeotime, China). The images of cell colony were captured.

### Annexin V-FITC/PI staining assay

The apoptotic ratio was measured with an apoptosis detection kit (Byeotime, China) according to the manufacturer’s instructions. Briefly, after treatment with the indicated concentrations of CHE for 24 hours, cells were collected by trypsinization and washed with PBS for Annexin V-FITC/PI staining. For each sample, 5×10^4^ cells were harvested and analyzed by flow cytometry (CytoFLEX, Beckman Coulter, Brea, CA, USA). Annexin V−PI−, Annexin V−PI+, Annexin V+PI−, Annexin V+PI+ staining represents viable cells, necrotic cells, early apoptotic cells, and late apoptosis cells being stained, respectively.

### Ubiquitination assay

HA-ubiquitin, HA-K48-ubiquitin, HA-K63-ubiquitin and Flag-PARP1 (wild-type or mutant) were co-transfected into HEK293T cells using PEI transfection reagent. After 48 h, cells were harvested and subjected to immunoprecipitation with an anti-Flag antibody to isolate PARP1, followed by immunoblotting with an anti-HA antibody to evaluate its ubiquitination. In selected experiments, transfected HEK293T cells were additionally treated with 12 or 24 mM sodium lactate for 24 h.

### *In vitro* lactylation assays

Purified recombinant protein PARP1 (10 μM) was mixed with 1 μM recombinant AlaRS in a 50 μL reaction buffer composed of 50 mM HEPES (pH 7.5), 25 mM KCl, 2 mM MgCl_2_, 5 mM sodium lactate, and 5 mM ATP. The mixture was incubated at 25°C for 2 h. Reactions were halted by adding 5×SDS loading buffer, and lactylation of PARP1 was assessed via western blot with an anti-pan-Klac antibody.

### LC-MS/MS analysis of the lactylated modification site

The sample was subjected to SDS-PAGE followed by Coomassie blue staining solution. The target bands were excised and peptides were extracted. Peptide samples were injected onto a custom-made C18-packed capillary column (20 cm length by 100 μm inner diameter, Dr. Maisch GmbH), and analyzed using an Orbitrap Exploris 480 mass spectrometer coupled with an EASY-nLC 1000 system (Thermo Fisher Scientific). The HPLC used a gradient of 5%-35% buffer B (0.1% formic acid in 80% acetonitrile) in buffer A (0.1% formic acid in water) over 20 min with a flow rate of 0.3 μL/min. In positive-ion mode, full-scan mass spectra were acquired from m/z 350 to 1,200 at a resolution of 60,000. Data-dependent MS/MS was performed on the 20 most intense ions at a resolution of 15,000 using higher-energy collisional dissociation with parameters set for an isolation window of 2.0 m/z, charge 2+, collision energy 30% and dynamic exclusion after two occasions within 20 s. Data were collected by Xcalibur installed on the mass spectrometer (Thermo Fisher Scientific), and MS/MS spectra were queried against the reverse, concatenated UniProt human FASTA database using Proteome Discoverer 2.4.

### Xenograft mouse model

Six-week-old female Balb/c nude mice were housed at the SPF animal facility. All experimental procedures and methods in mice were conducted in accordance with the institutional guidelines approved by the Animal Care and Use Committee (IACUC) of Shanghai Jiao Tong University (Issue No. KT2022608). Wild-type S26 cells were injected subcutaneously at a dose of 3 × 10^5^ cells per mouse, and CHE (0.1 mg/kg, dissolved in solvent including 1% DMSO, 30% PEG300 and 8% Tween 80) or solvent was injected into the tumor every 2 days (n = 5/group). After 25 days of continuous observation, the mice were sacrificed, and the weights of xenografted tumors and mice were measured.

### Human HNSCC samples

The use of human specimens in this study was approved by the Ethics Committee of Shenzhen Third People’s Hospital (Approval No. 2021-055). HNSCC tissue samples were obtained from biopsy-confirmed patients. Matched non-tumor control tissues were collected from histologically normal adjacent regions. All participants provided written informed consent prior to inclusion.

### Tissue collection and histological analysis

Xenograft tumors and patients’ tissue were fixed, sectioned, and stained according to the manufacturer’s instructions (ZSGB-BIO, Cat#PV-6000, ZLI-9017). IHC staining of Mortalin, PARP1, and AlaRS were quantitatively analyzed using Image J, and the average optical density (AOD) was used for statistical analysis. AOD= integrated optical density (IOD)/area of positive staining in each IHC staining image.

### RNA-seq profiling and analysis

Total RNAs were isolated from AlaRS^KD^, CHE treatment, and Control S26 cells (n = 3/group) to generate RNA libraries. The Analysis and profile were performed at Huada Sequencing Corporation. Genes with adjusted p value < 0.05 or the absolute log2-fold change >0 were considered as differentially expressed genes (DEGs). The obtained DEGs were further used to draw the volcano map and heatmap.

### qRT-PCR

Total RNA extraction was performed using TRIzol (Vazyme). Reverse transcriptase (#639538, TAKARA) along with oligo-dT primers was applied for reverse transcription. SYBR green qPCR Mix kit (Aaccurate biology) was used for the qRT-PCR process. The transcription level of GAPDH served as the endogenous control. All data were normalized and calculated via the comparative critical threshold cycle 2^-ΔΔCt^ method. The primer sequences are listed in Table S6.

### Western blot assay

Total protein was extracted from cells or tissues by the RIPA buffer (50 mM Tris-HCl pH 7.4, 150 mM NaCl, 1% NP40, 1 mM EDTA), supplemented with protease and phosphatase inhibitor cocktail (Roche)). After SDS-PAGE, the protein on the gel was transferred to a PVDF membrane (Bio-Rad, Hercules, CA, USA) and then blocked with 5% non-fat dry milk in TBST for 1 hour. Then, the membranes were incubated with indicated primary antibodies and corresponding HRP-conjugated secondary antibodies. Signals were detected by Bio-Rad ChemiDoc system (Bio-Rad, Shanghai, China) and analyzed using ImageJ software. The following antibodies were used: AlaRS antibody (Abcam, Cambridge, UK, 1:3000), PARP1 (Santa cruz, 1:100), p21 (ABclonal, Wuhan, China, 1:1000), p53 (ABclonal, Wuhan, China, 1:1000), P-RB (Cell Signaling Technology, Boston, MA, USA, 1:3000), Mortalin (ABclonal, Wuhan, China, 1:1000), GAPDH (Ray antibody biotech, Beijing, 1:5000), or beta-actin (Cell Signaling Technology, Boston, MA, USA, 1:1000).

### Molecular docking

Autodock software was used for molecular docking. The structure of the AlaRS^1–84^ and ligand CHE were prepared before docking. The macromolecule receptor was modified by removing water molecules, adding polar hydrogen atoms, and then the semiflexible docking method was used to generate the model. Molecular visualization and analysis were performed using Pymol version 2.5.4 (Schrödinger, LLC).

### Co-Immunoprecipitation (Co-IP)

Whole-cell lysates (400 μg) were treated overnight with 3 μg of anti-Flag antibody in RIPA buffer. Next, Protein A/G agarose beads (#sc-2003, Santa Cruz Biotechnology) were applied to the lysates and the samples were further treated for 4-6 hours with rotation. Samples were washed three times with RIPA buffer. Subsequently, the immunoprecipitations were resuspended into 2×SDS-loading buffer. The samples were then boiled for 10 min and detected using western blot.

### Glutathione S-transferase (GST) pull-down assay

AlaRS full-length and AlaRS/Mortalin fragments fused with 6×His were purified with Ni-NTA beads, followed by chromatography on a HiTrap Q HP column. Mortalin or PARP1 fragments fused with GST were purified with Glutathione Beads (#SA008100, Smart-Lifesciences). For GST pull-down assays, 200 μg of each GST-fused protein was mixed with an equal amount of 6×His-tagged fusion protein in binding buffer (50 mM Tris-HCl pH 7.5, 20 mM NaCl, 5% glycerol, 5 mM DTT with 0.1% BSA) at 4°C overnight. The mixture was next incubated with 20 μL glutathione beads at 4°C with constant rotation for 2 hours. Subsequently, the beads/protein complex was washed five times, and eluted with 20 μL 2×SDS sample buffer. The samples were then subjected to western blot analysis with anti-GST ((Proteintech, Chicago, IL, USA, 1:3000), anti-His (#YM2096 Immunoway, 1:3000) antibodies.

### Immunoprecipitation LC-MS/MS (IP-MS)

After immunoprecipitation, LC-MS/MS was conducted to identify the potential interacting protein of AlaRS in HEK293T cells. The immunocomplexes were eluted by SDT buffer (4% SDS, 100 mM Tris-HCl, 1mM DTT, pH 7.6), followed by boiling for 10 min. LC-MS/MS experiments were conducted by Applied Protein technology (Shanghai).

### Statistics and reproducibility

Statistical analyses were performed using SPSS 25 or GraphPad Prism v.10 software. Numerical variables are presented as mean ± standard deviation (SD) or standard error of the mean (SEM). Differences between two groups were compared using two-sided Student’s t-tests or Wilcoxon rank-sum tests, as appropriate. For comparisons among multiple groups, two-sided Analysis of Variance (ANOVA) was employed. The Spearman correlation coefficient was used to analyze correlations between numerical and categorical variables. Survival curves were generated using the Kaplan-Meier method and compared via the log-rank test. Statistical significance was set at p < 0.05 and denoted by asterisks: *p < 0.05, **p < 0.01, ***p < 0.001, ****p < 0.0001. Non-significant p-values were indicated as “ns”.

For cell-based quantifications, cells were randomly selected according to pre-defined inclusion criteria relevant to the experimental objective. In mouse experiments, age- and sex-matched animals were randomly assigned to experimental groups to ensure balanced distribution. Data collection and analysis were not performed blind to the experimental conditions, and no samples or animals were excluded from the analysis. All experiments were independently replicated, and comparable results were obtained across replicates. Representative images for western blotting and microscopy were derived from at least two independent experiments.

### Large language models usage statement

During the preparation of this work, Gemini (Google) was used to assist with language editing, formatting, and polishing of the manuscript. After using this tool, the authors reviewed and edited the text as needed and take full responsibility for the final content of the publication.

## Data availability

The RNA-seq data generated in this study have been deposited in the NCBI Sequence Read Archive (SRA) under BioProject accession SRP524184. The mass spectrometry proteomics data have been deposited to the ProteomeXchange Consortium via the iProX partner repository with the dataset identifier PXD055806. All other relevant data supporting the findings of this study are available from the corresponding author upon reasonable request.

## Acknowledgements

We acknowledge Professor Na Yang from Institute of Microbiology, Chinese Academy of Sciences, for kindly providing the human pET-28a-sumo-SIRT1 and pRSFDuet-sumo-SIRT2 plasmids. S18 and S26 cells were gift from Professor Hua Zhang at Medical School of Sun Yat-sen University. We thank Professor Caijun Sun at School of Public Health (Shenzhen) of Shenzhen Campus of Sun Yat-sen University for providing Lenti-virus system. We thank Chenchen Zhu at Medical School of Sun Yat-sen University for technical support. This work was supported by the National Natural Science Foundation of China (Grant No. 32271314, China), the Shenzhen Science and Technology Innovation Commission (Grant No. ZDSYS20230626091203007 and JCYJ20240813151452068, China).

## Authors Contributions

J.Z., X.M., Q.Q., Q.Z., Y.L., R.Q., Y.Y., C.L., and G.L. performed the experiments; J.Z., X.M. and L.S. designed the experiments; X.L., Y.B., Z.Z., and J.G. were involved in methodology design; J.Z., X.M., S.Z., L.W., B.M. and L.S. analyzed the data; Z.Z., J.Z., J.G., L.W., B.M., P.S., and L.S. reviewed and revised the draft. All authors read the final version of the manuscript and approved the final manuscript.

---

**Figure S1.**
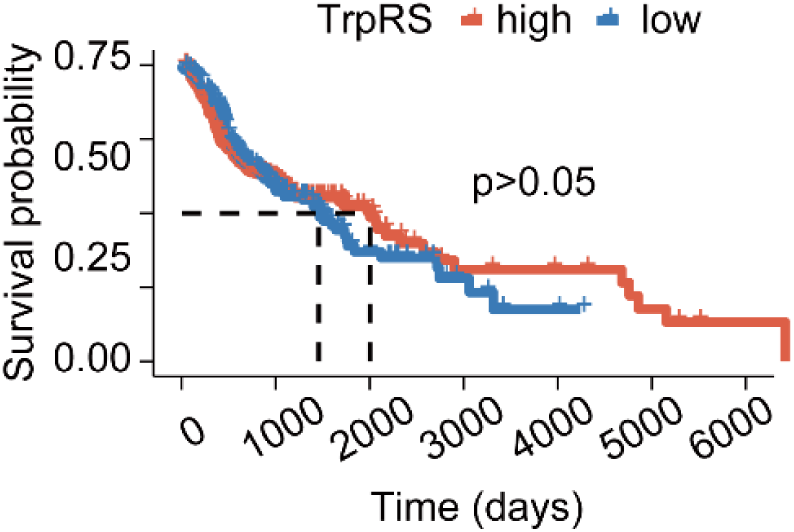
Relationship between TrpRS expression and survival in HNSCC patients. The Kaplan-Meier survival curve depicting overall survival relative to TrpRS expression in diagnosed HNSCC patients (Log-rank test, p > 0.05).

**Figure S2.**
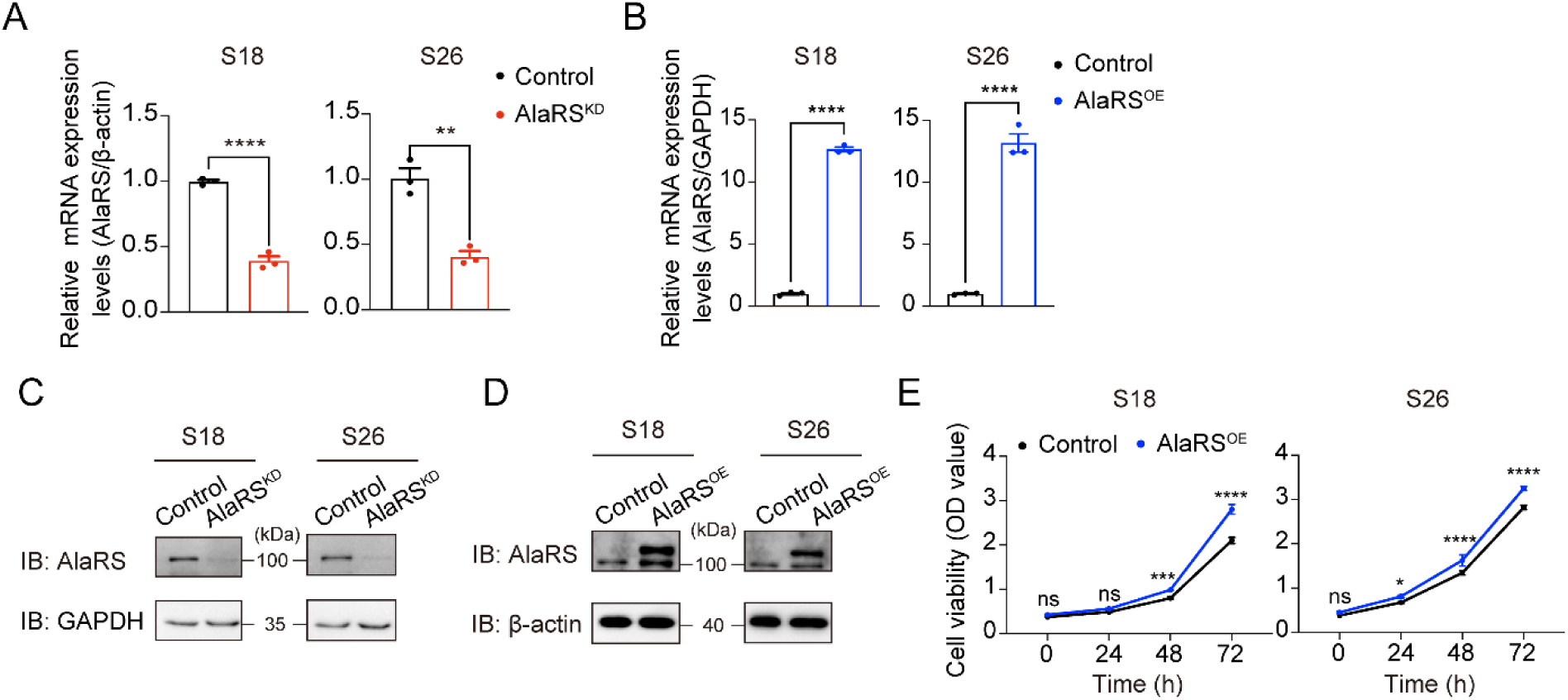
AlaRS knockdown and overexpression in S18 and S26 cells. **A-B)** qRT-PCR analysis of *AlaRS* mRNA levels after knockdown (shRNA) or overexpression (PLVX-AlaRS-GFP) in S18 and S26 cells (n = 3). ** p < 0.01, **** p < 0.0001. **C-D)** Western blot analysis of AlaRS protein levels in S18 and S26 cells after AlaRS knockdown **(C)** or overexpression **(D)**. **E)** Cell viability assessed by CCK-8 assay in S18 and S26 cells overexpressing AlaRS (n = 3). ns, not significant, *p < 0.05, ***p < 0.001, ****p < 0.0001.

**Figure S3.**
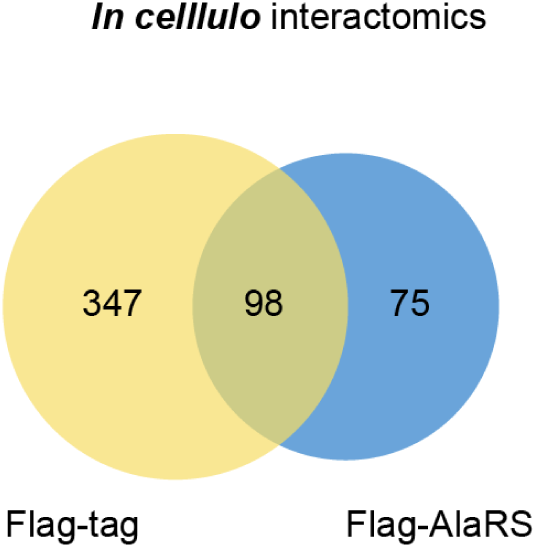
Venn diagram of proteins identified by IP-MS. Venn diagram showing the distribution and overlap of proteins identified through IP-MS analysis.

**Figure S4.**
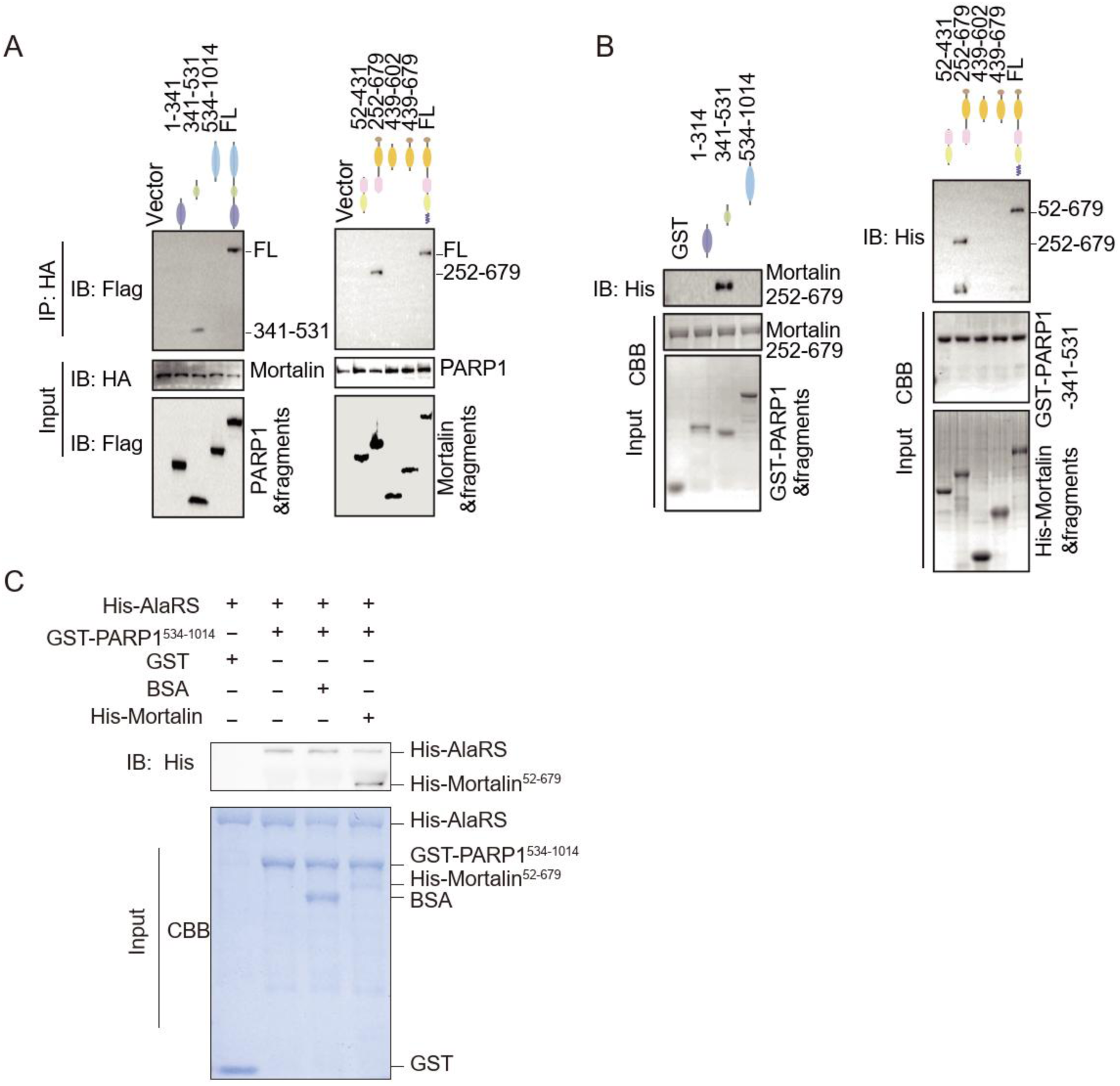
PARP1 directly interacts with Mortalin. **A)** Co-IP assays with truncated PARP1 and Mortalin constructs to map interacting regions. **B)** GST pull-down assays with truncated proteins to define the binding domains of PARP1 and Mortalin. **C)** *In vitro* pull-down assay assessing the interaction between purified AlaRS and PARP1 in the presence of recombinant Mortalin or BSA.

**Figure S5.**
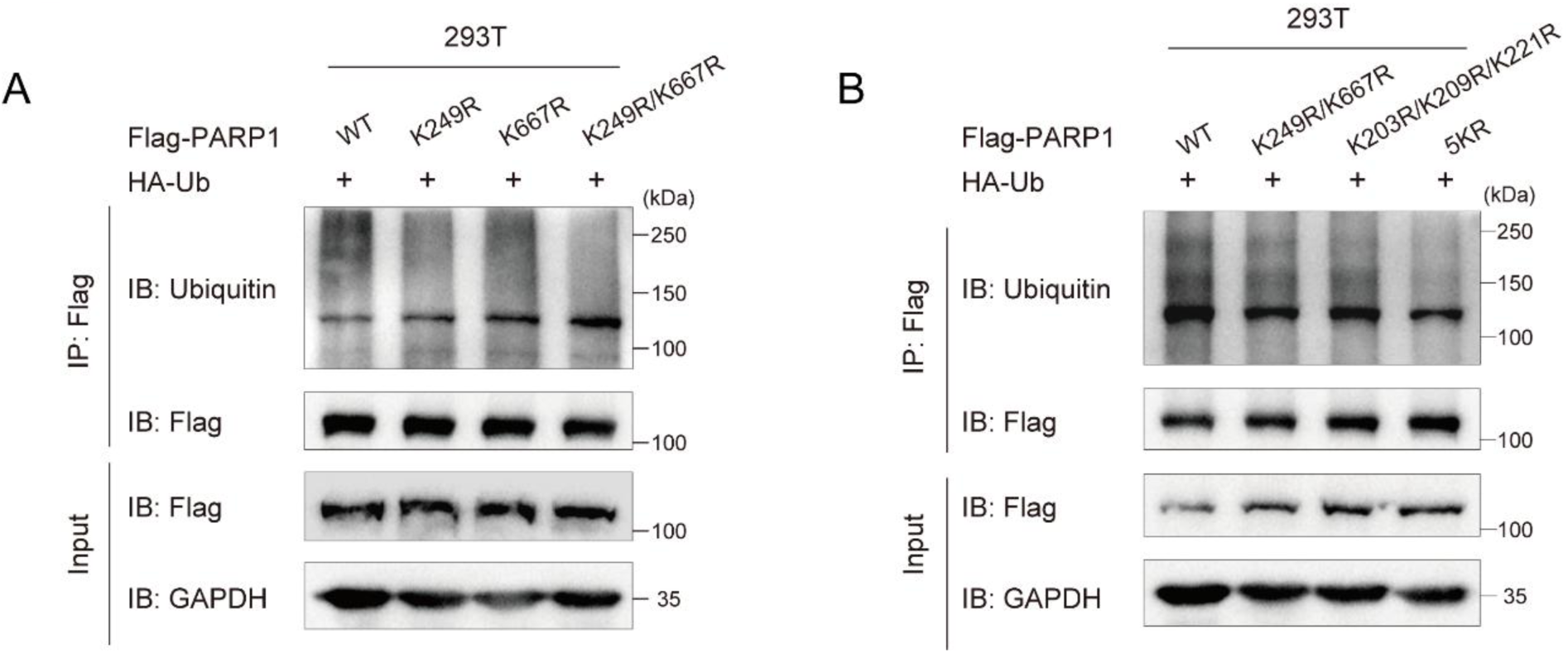
Ubiquitination analysis of site-directed PARP1 mutants. **A-B)** Ubiquitination of PARP1 in HEK293T cells co-expressing HA-Ub and Flag-PARP1 (WT or indicated lysine mutants). 5KR: K203R/K209R/K221R/K249R/K667R.

**Figure S6.**
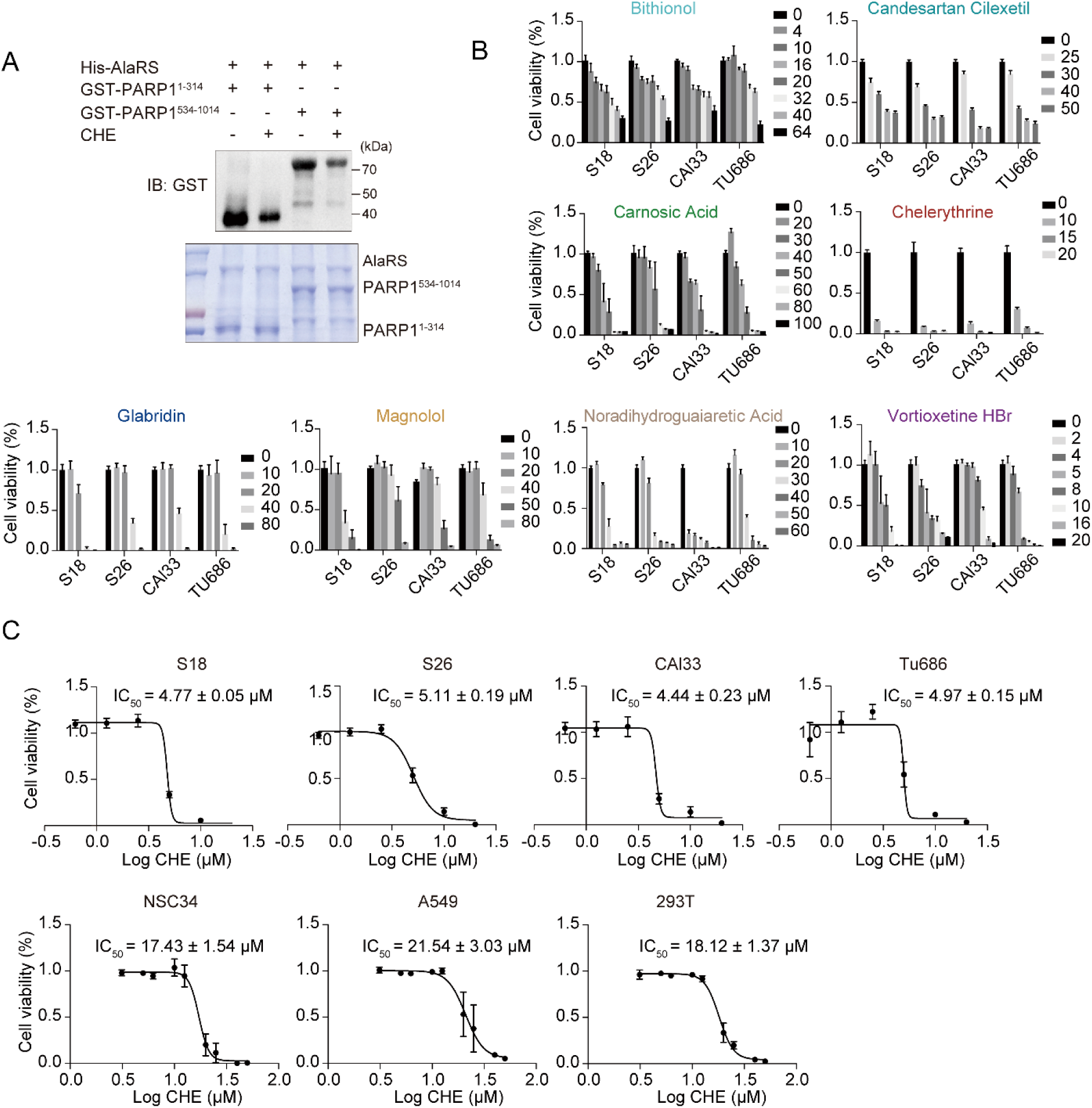
Validation of small-molecule-induced inhibition of cell viability. A) Western blot analysis of AlaRS-mediated PARP1 lactylation *in vitro* in the presence or absence of CHE. B) Inhibitory activity of eight candidate compounds against four HNSCC cell lines, assessed by CCK-8 assay (n = 3). C) Dose-response analysis to determine the IC_50_ of CHE in a panel of cell lines, evaluated by CCK-8 assay (n = 3).

**Figure S7.**
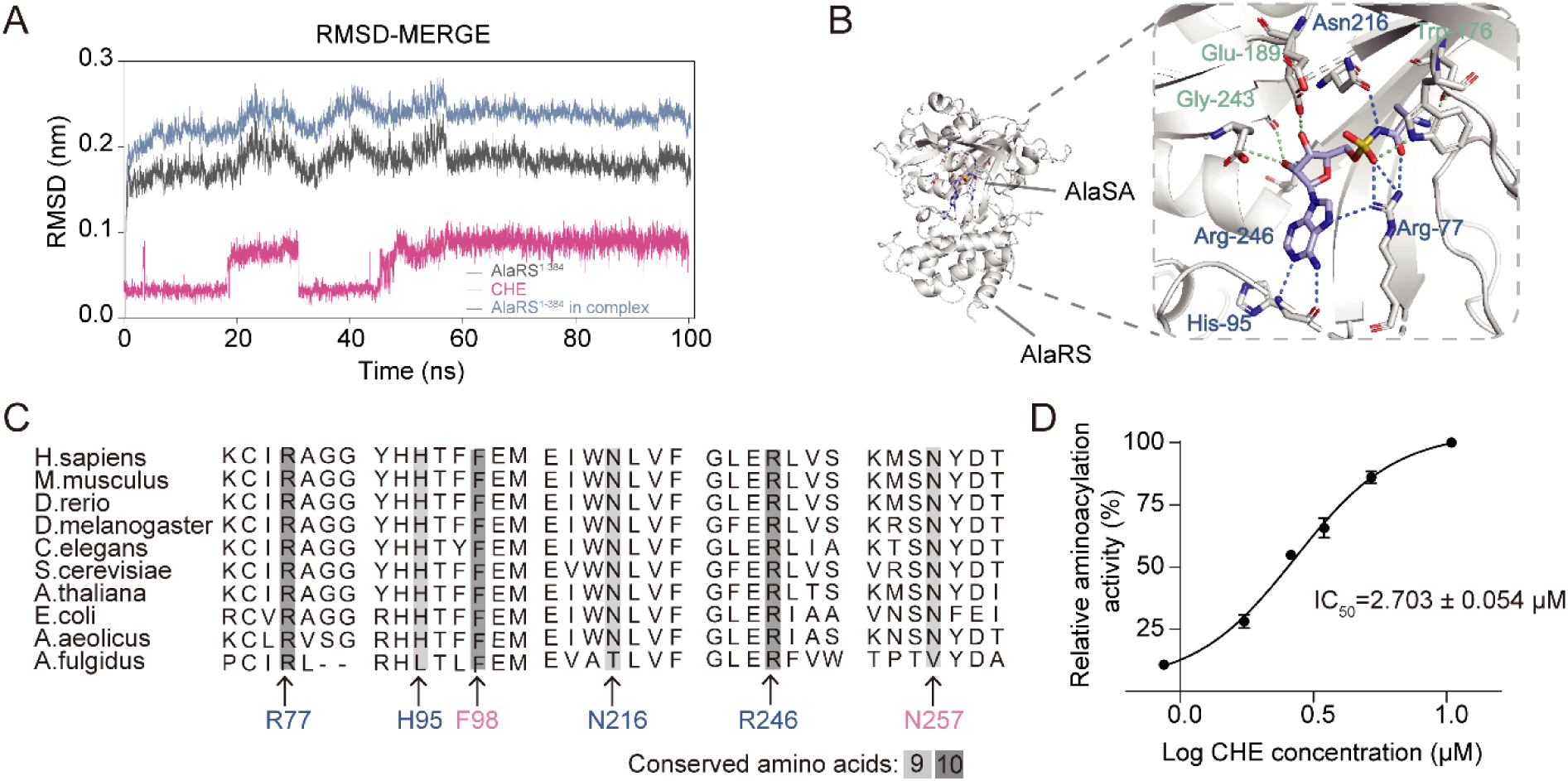
Structural characterization of the AlaRS-CHE interaction. **A)** Root-mean-square deviation (RMSD) of CHE, the AlaRS^1–84^ fragment, and their complex during molecular dynamics simulations. **B)** Binding mode of AlaSA (an Ala-AMP analog) to human AlaRS^1–84^ (PDB: 4XEM), shown in colored stick representation. **C)** Evolutionary conservation analysis of potentially CHE-interacting residues across eukaryotic, bacterial, and archaeal AlaRS orthologs (aligned by Clustal Omega). **D)** Dose-dependent inhibition of AlaRS aminoacylation activity by CHE, measured by ATP consumption (n = 3).

**Figure S8.**
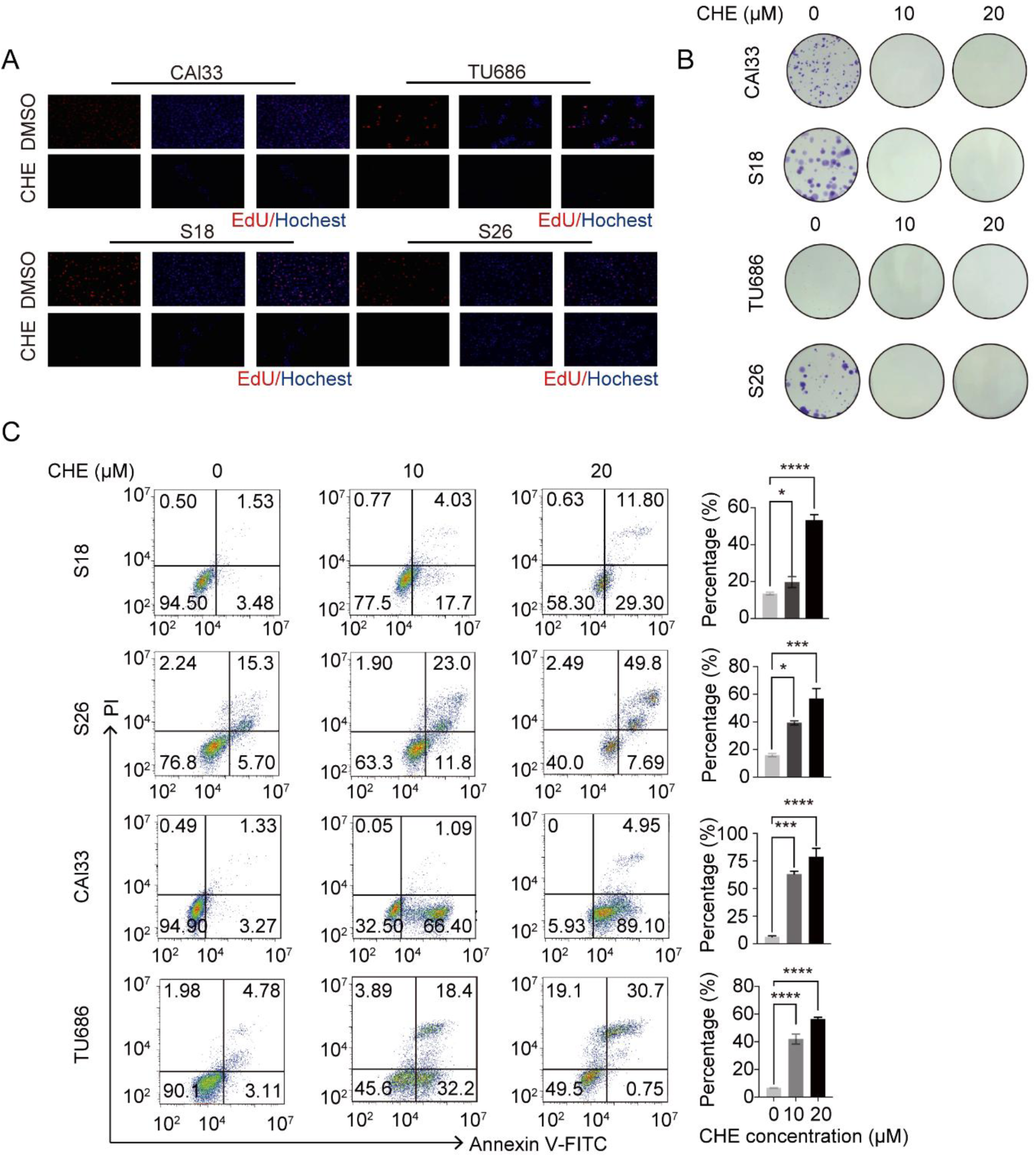
Functional assessment of CHE in HNSCC cells. **A)** Representative images of EdU staining images of S18, S26, CAl33 and TU686 cells treated with DMSO or 10 μM CHE for 24 h. Scale bars, 40 µm. **B)** Colony formation assay of HNSCC cells treated with CHE for 24 h. Colonies were stained with crystal violet and representative images are shown. **C)** Apoptosis assessed by Annexin V-FITC/PI staining after 6 h of CHE treatment at indicated concentrations. *p < 0.05, ***p < 0.001, ****p < 0.0001.

**Figure S9.**
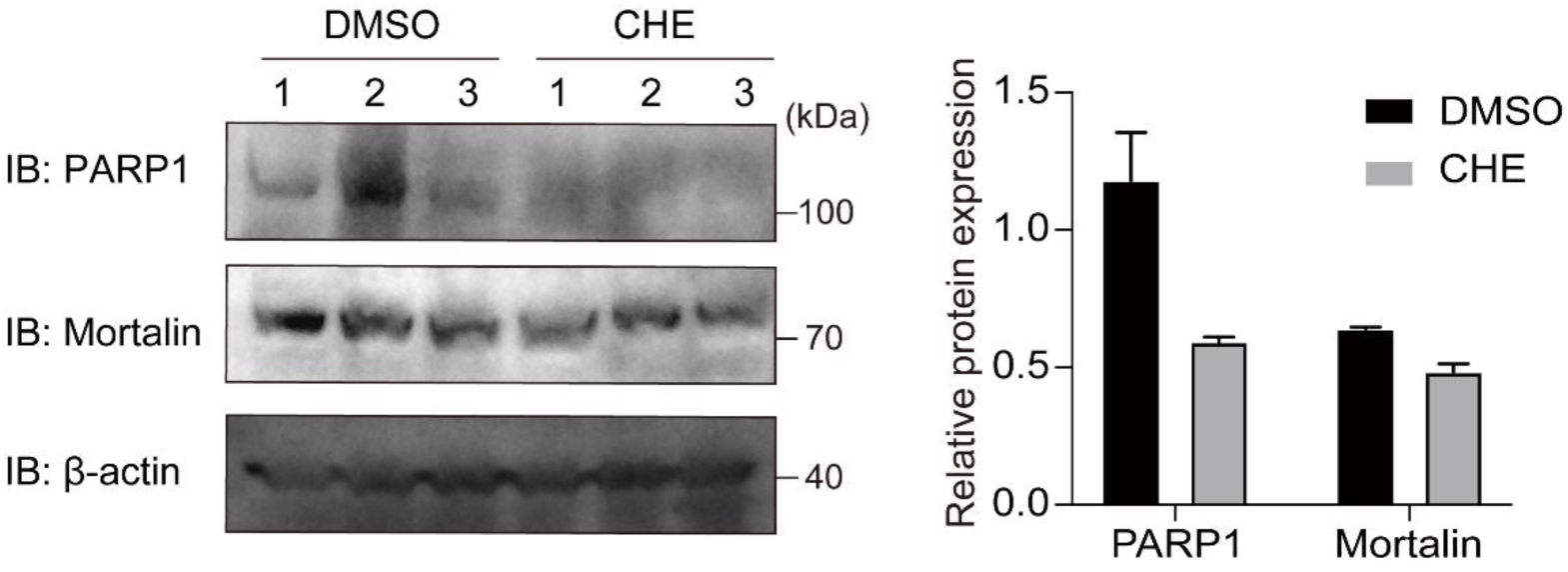
Effects of CHE treatment on PARP1 and Mortalin protein levels in HNSCC xenograft tumors. Western blot analysis and quantification of PARP1 and Mortalin protein levels in tumor tissues from the xenograft model after CHE treatment (n = 3).

**Table S1.** Characteristics of 15 HNSCC patients.

| Factors | Number | Percentage (%) |
| --- | --- | --- |
| Age |  |  |
| >=60 | 8 | 53.3 |
| <60 | 7 | 46.7 |
| Gender |  |  |
| Male | 13 | 86.7 |
| Female | 2 | 13.3 |
| T stage |  |  |
| T1 | 2 | 13.3 |
| T2 | 3 | 20 |
| T3 | 9 | 60 |
| T4 | 1 | 6.7 |
| Lymph node involvement |  |  |
| N0 | 9 | 60 |
| N1 | 4 | 26.7 |
| N2 | 2 | 13.3 |
| N3 | 0 | 0 |
| Tumor site |  |  |
| Larynx | 6 | 40 |
| Hypopharynx | 4 | 26.7 |
| Oral cavity | 5 | 33.4 |
| Smoking |  |  |
| Yes | 12 | 80 |
| No | 3 | 20 |
| Drinking |  |  |
| Yes | 7 | 46.7 |
| No | 8 | 53.3 |

**Table S2.**
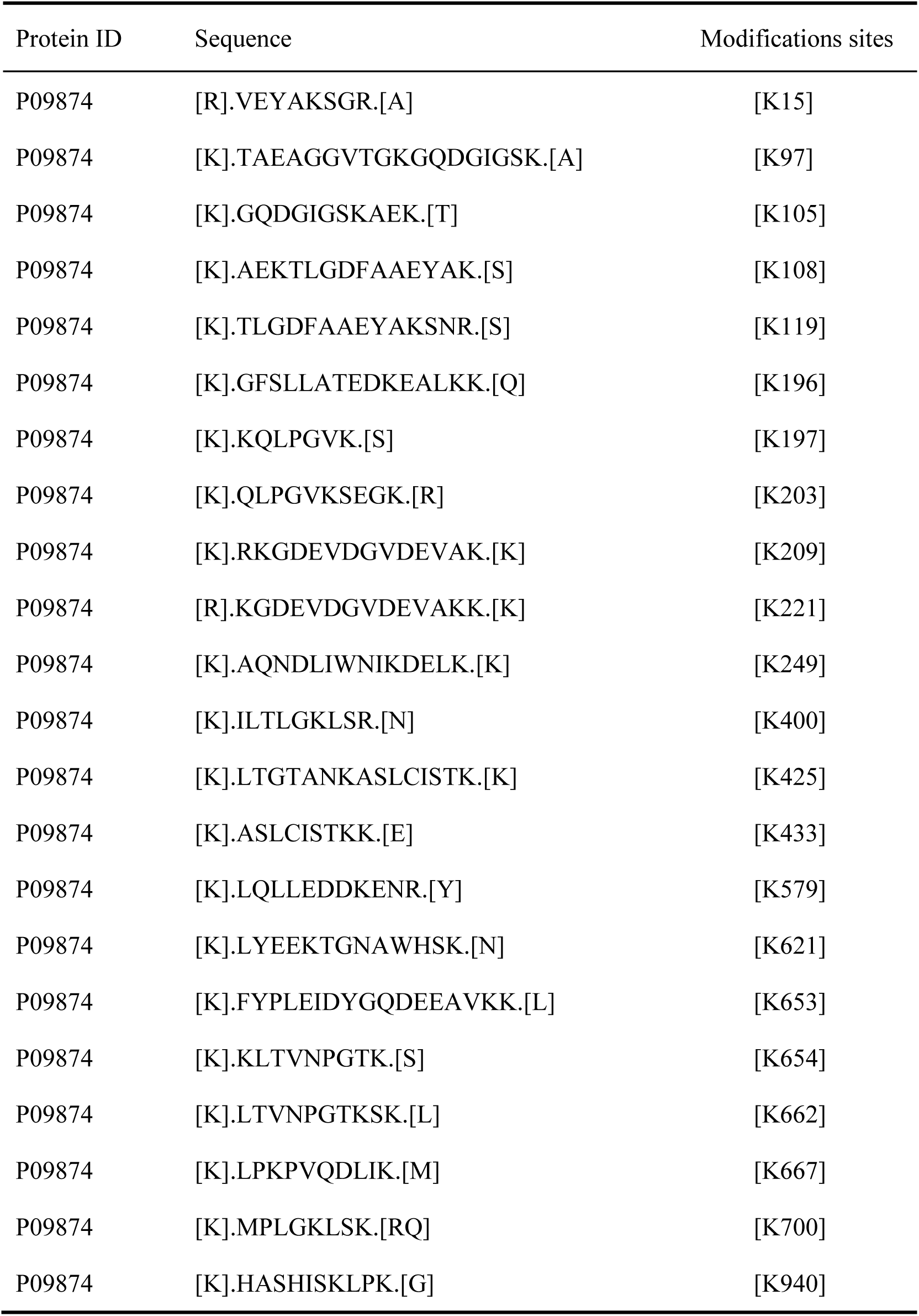
Lactylation sites information of PARP1.

**Table S3.** BLI values of 14 hits interacting with AlaRS.

| Compounds | K<sub>D</sub> (μM) | K<sub>on</sub> (1/Ms)×10<sup>3</sup> | K<sub>dis</sub> (1/s)×10<sup>-2</sup> |
| --- | --- | --- | --- |
| A | 1.79±0.26 | 35.27±8.85 | 6.16±0.74 |
| B | 11.75±0.18 | 5.74±0.55 | 6.75±0.75 |
| C | 20.08±9.39 | 1.73±1.02 | 7.27±4.47 |
| D | - | - | - |
| E | 10.39±4.49 | 49.38±37.51 | 22.29±1.84 |
| F | - | - | - |
| G | - | - | - |
| H | - | - | - |
| I | 15.92±1.82 | 16.50±0.01 | 26.27±3.03 |
| J | 39.71±7.24 | 25.59±1.58 | 102.20±24.79 |
| K | - | - | - |
| L | - | - | - |
| M | 13.26±3.56 | 6.14±5.05 | 11.42±6.14 |
| N | 48.22±3.68 | 25.27±17.82 | 54.93±30.06 |

Compounds: A, Bithionol; B, Candesartan Cilexetil; C, Carnosic Acid; D, Ceftributen Dihydrate; E, Chelerythrine; F, Cynarin; G, Fupentixol Dihydrochloride; H, Ginsenoside Rb1; I, Glabridin; J, Magnolol; K, Methylcobalamin; L, Momordin Ic; M, Nordihydroguaiaretic Acid; N, Vortioxetine HBr

**Table S4.**
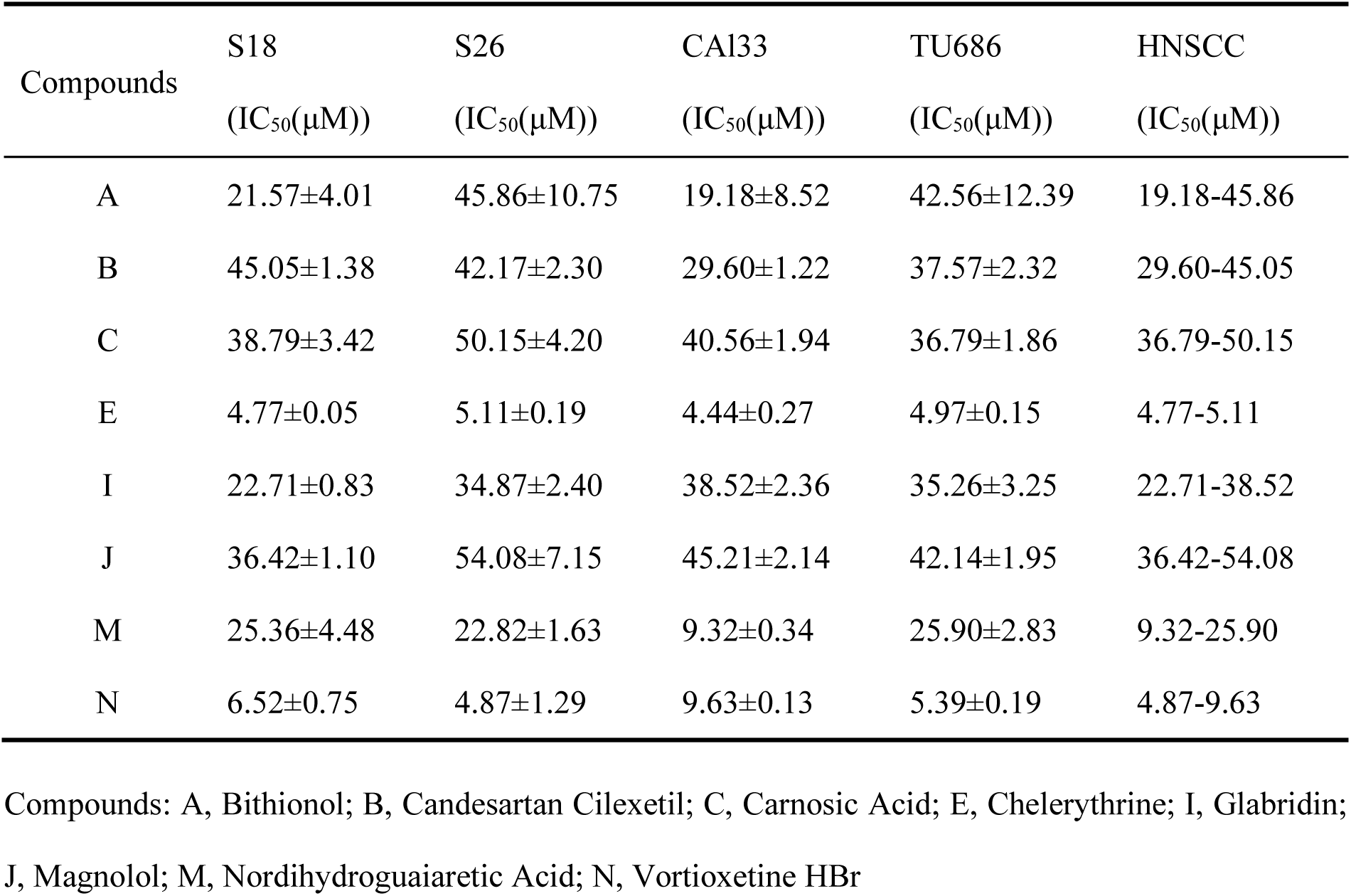
IC_50_ of 8 hits in HNSCC cell lines.

**Table S5.**
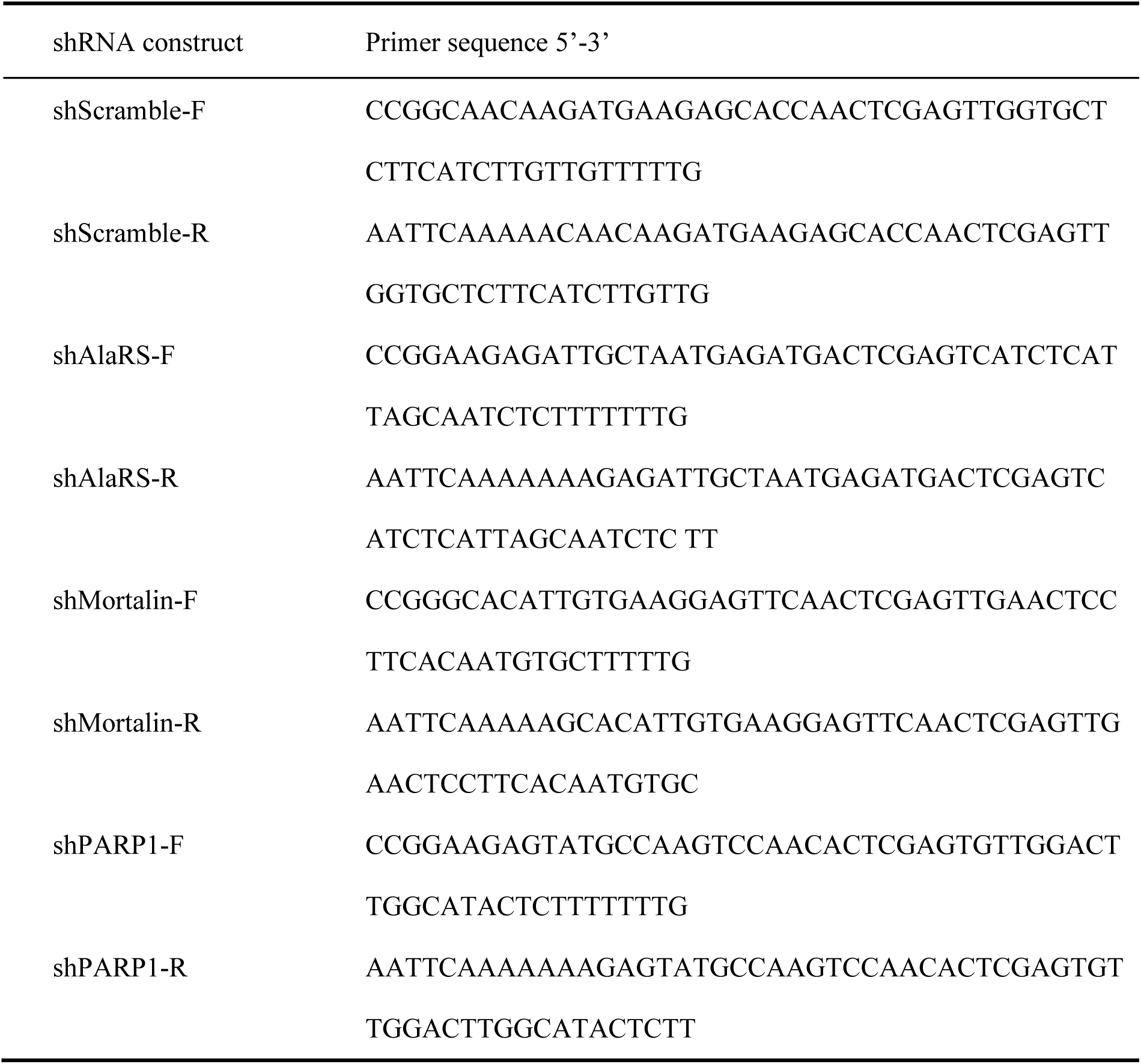
Primers used for shRNA construct.

**Table S6.**
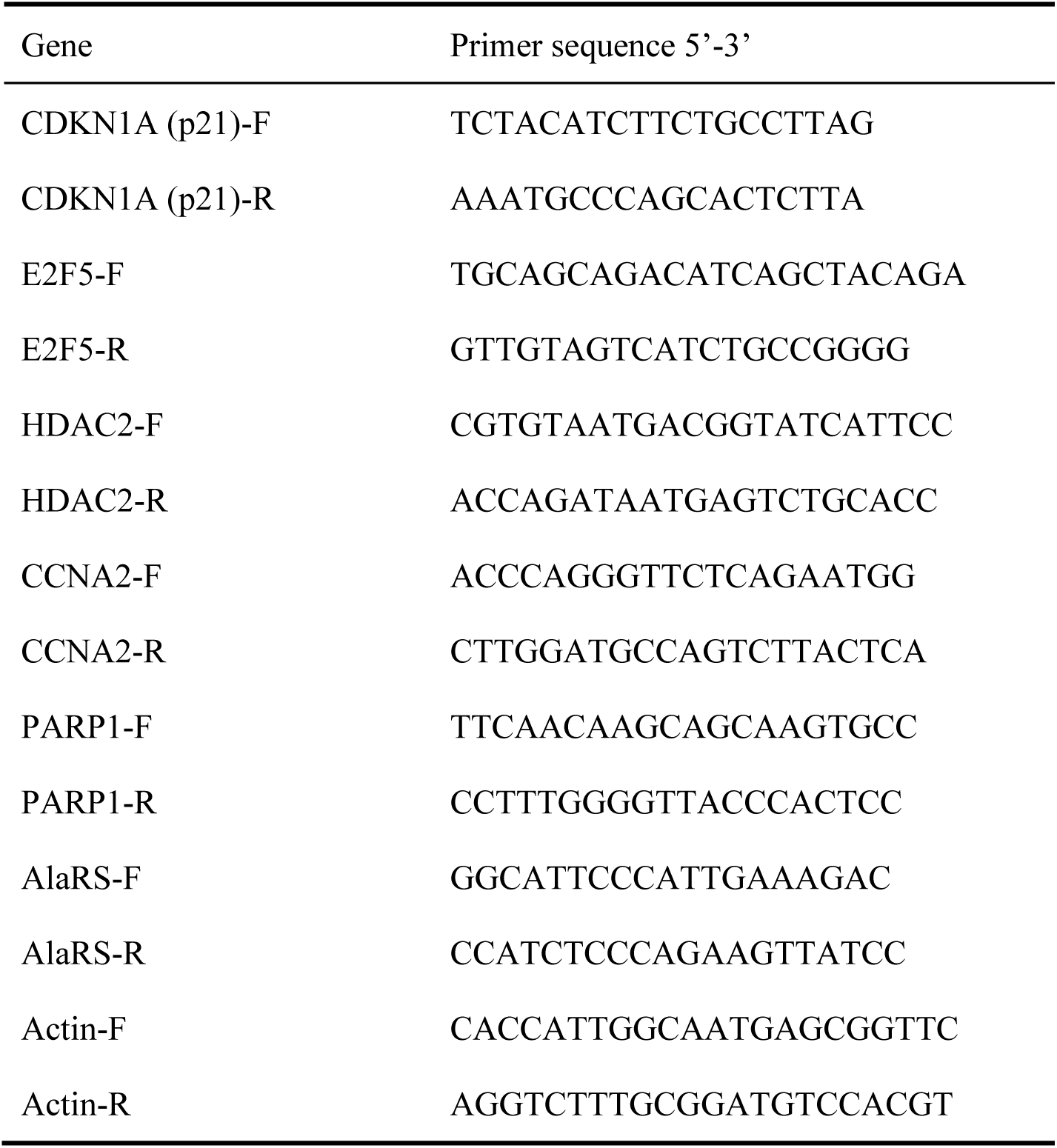
Primers used for qRT-PCR.

